# *In silico* analysis and cluster validation of potential breast cancer nsSNPs of serological tumor marker CA27.29

**DOI:** 10.1101/2021.01.19.427371

**Authors:** Jyoti Lakhani, Dharmesh Harwani

**Author notes:** **Address Correspondence to**, Dharmesh Harwani (Ph.D.), Kanav Bhawan, Academic Block 1 Department of Microbiology Maharaja Ganga Singh University Bikaner-334004, Rajasthan, India.

## Abstract

**Background & Objectives:** CA27.29 is a breast cancer-associated glycoprotein. Many genetic variations caused by non-synonymous nucleotide polymorphisms (nsSNPs) are known to affect the functionality of the CA27.29 protein.

**Methods:** In the present manuscript, an *in silico* analysis of the genetic variations in CA27.29 was done to observe functional nsSNPs that possibly alter its stability. Among 2205 SNPs identified from the publically available SNP database (dbSNP), 213 (9.66%) synonymous SNPs, 24 (1.09%) non-synonymous SNPs, and 1351 (61.27%) noncoding intronic SNPs were observed. The function predictability tools SIFT, Provean and Polyphen2 were used to uncover variations in the analyzed nsSNPs and their functionality.

**Results:** A total of 16, 20 and 10 non-synonymous SNPs (nsSNPs) were predicted to be damaging or deleterious by SIFT, Polyphen, and Provean tools respectively. Intriguingly, 9 nsSNPs were predicted to be damaging by all the three tools used while 4 nsSNPs were predicted to be damaging by SIFT and Polyphen tools. The substitutions C/G->T and G->A/T were observed to be dominant in the analyzed nsSNPs that probably have damaging role to CA27.29 glycoprotein. Moreover, the validation of results using ClinVar tool revealed that all the analyzed possibly, probably and highly damaging nsSNPs are yet to be reported and studied. Besides this, we found Global minor allele frequency (Global MAF) for only 11 nsSNPs, the values of which were observed to be <0.1% that further confirmed the novelty of the analyzed variants.

**Interpretation & Conclusions:** Among the analyzed nsSNPs, 3 nsSNPs rs145691584, rs148332231 and rs191544901 were found to be located in 3’UTR region of CA27.29 gene that were assumed to have the possible functional roles in altering the protein stability. The present study is useful to gain useful insights into the genetic variations in nsSNPs that may playing a critical role in determining the susceptibility to breast cancer.

## INTRODUCTION

About one in four of all new cancer cases (24.2%) diagnosed worldwide in women is breast cancer^1^. According to the press release from the International Agency for Research on Cancer (IARC), breast cancer is the leading cause of cancer death in women (15.0%), followed by lung cancer (13.8%) and colorectal cancer (9.5%)^2^. The disease caused 627000 deaths in 2017-18 which is 6.6% of the total mortality rate that year^3^. It is possible to treat early-stage breast carcinoma successfully but once this melanoma enters in the advanced or metastatic stage, it will become incurable. Consequently, it is important to recognize the symptoms of metastasis in breast cancer patients as early as possible. The truquant BR radioimmunoassay test or CA27.29 test is the most commonly used metastatic breast cancer detection test that helps medical practitioners to diagnose the reappearance of carcinoma cells after its successful treatment. The normal measurement of CA27.29 antigen in the blood is less than 40U/ml. The higher the measurement of CA27.29 antigen, the greater the likelihood of the presence of breast cancer cells in the body. This test is based on the breast cancer-associated antigen known as CA27.29 (MUC-1; *mucin* gene product), a special glycoprotein present on the breast cancer epithelial cells. Breast cancer cells shed copies of CA27.29 protein into the bloodstream in case of metastatic breast cancer or in advanced stages of breast cancer^4,5^ [4] [5]. The other tumor markers are CA15-3, carcinoembryonic antigen CEA, and HER2 extracellular domain (ECD)^6^.

The guidelines of the American Society of Clinical Oncology (ASCO) recommends the use of breast cancer tumor markers such as CA15.3, CA27.29, and CEA in the breast cancer diagnosis^6,7^ [6][7]. Among these, CA15.3 and CA27.29 tumor markers have received also US Food and Drug Administration (FDA) approval in 1996 for monitoring of the disease recurrence or metastasis in patients with stage II/III breast cancer^8^. However, ASCO also recommends that these tumor markers should not be used alone for the surveillance of disease during active therapy and be used in conjunction with diagnostic imaging, history, and physical examination. Conversely, some studies recognize MUC-1 tumor markers CA15.3 and CA27.29 as independent predictors for the primary diagnosis of the disease in addition to the traditional markers such as tumor size and nodal status^9^. The studies also indicate that after primary treatment, the level of these markers can predict the disease recurrence about six months in advance than the other available methods^10,11^. However, according to the current guidelines, the tumor markers CA15-3 and CA27.29 are not recommended as the prognostic markers for routine clinical use. Despite these limitations, these tumor markers are widely used in disease surveillance and treatment monitoring^12^. In a study, medical practitioners attempted a trial to evaluate the role of the tumor marker CA27.29 after surgery before taxane-based adjuvant chemotherapy in 2669 primary breast cancer patients and they found that CA27.29 levels were above the cut-off level in 8% of the analyzed patients. The study provided information on the prognostic relevance of CA27.29 before the systemic treatment and demonstrated the role of this tumor marker in adjuvant chemotherapy, endocrine, bis-phosphonate, and treatment monitoring^12^. Thus the role of CA27.29 has been observed to be inestimably useful in identifying the response of the patients to the breast cancer treatment and re-occurrence of the disease after treatment. Importantly, other than breast cancer, lung cancer, liver cancer, pancreatic cancer, colon cancer, ovarian cancer, prostate cancer also show elevated levels of CA27.29 antigen^13^.

The functionality of a gene cannot be fully understood without knowledge of the potential variability within a gene^14^. Genetic variants can have a major impact on gene function. Single nucleotide polymorphisms (SNPs) are the most common form of human genetic variations^42^. It has been estimated that one SNP exists every 290 base-pairs in the human genome^15^. Evidence showed that a wide range of human diseases including cancer can be triggered through SNPs^16,17^. SNPs might also affect the pharmacokinetics and pharmacodynamics of certain drugs in cancer therapy^18^. There are two groups of SNPs: synonymous (sSNP) and non-synonymous SNPs (nsSNP), the latter one results in changes of the translated amino acid sequence^19^. With the increasing number of known human nsSNPs, the studies pertaining about the identification of a subset of nsSNPs that affect protein function/s have also increased. Various types of features can be used to predict the functional impact of nsSNPs such as physical and chemical properties of the affected amino acids, structural properties of the encoded protein, and evolutionary properties. Sequence alignment of homologous proteins^20^. SIFT (Sorting Intolerant from Tolerant)^21^, PROVEAN (Protein Variation Effect Analyzer)^22^ and PolyPhen-2 (Polymorphism Phenotyping v2)^23^ are some reliable computational prediction tools that consider these features to predict whether a given nsSNP has a functional impact or not. To understand the biological function and regulation of CA27.29 and its potential for its use as a marker, in the present research, an *in silico* analysis of the genetic variations in CA27.29 gene was performed to identify possible functional nsSNPs. The study involved, CA27.29 associated 2205 SNPs that were downloaded from the NCBI dbSNP database. These nsSNPs were investigated and validated using various computational tools to assess their functional impact on CA27.29. The observations from the present study led to the identification of a small number of nsSNPs that presumed to affect the CA27.29 protein function adversely. The present study provides more insights to mark the functional nsSNPS that mediate the genetic variations in CA27.29 protein. The analyzed variations may perhaps alter the functionality of the protein and play a critical role in determining susceptibility to breast cancer.

## MATERIAL AND METHODS

### SNP database

The dbSNP database, the largest repository of SNPs with over 140 million submitted variations^24^, hosted by the National Center for Biotechnology Information (NCBI)^25^ was used for SNP mining.

### Prediction of Functionally Important nsSNPs

#### SIFT

The prediction tool SIFT^26,27,28^ comprehends the sequence homologies of SNPs and then evaluates the functional impact. The prediction of SIFT tool is based on the degree of conservation of each amino acid residue in the query sequence. SIFT scans the dataset of functionally related protein sequences to evaluate the degree of conservation. This is done by searching the protein databases, UniProt and TrEMBL, using PSI-BLAST algorithm. Here, in the first step, the identified sequences are aligned with the query sequences for the function prediction^19^. Thereafter, in the second step, each substitution at each position in the aligned sequences is assessed and the probability of each substitution is calculated and recorded in a probability matrix. The calculated probability of sequence substitution is called the SIFT Score. A sequence substitution is considered to be tolerated by the SIFT tool if the SIFT Score is above 0.05. The sequence substitution with SIFT Score under 0.05 is considered to be deleterious. SIFT presumes that the sequence substitution at the highly conserved position cannot be tolerated than the sequence substitution at poorly conserved positions^29^.

#### PROVEAN

PROVEAN^22,30,31^ uses an alignment approach to evaluate the functional impact of SNPs similar to the SIFT tool. It consists of two main steps. In step 1, BLASTP is performed and similar sequences are collected from NCBI NR Protein database. The sequences with 80% or more similarity are clustered and termed as supporting sequences. In the second step, the delta score is calculated for each cluster of supporting sequences using the BLOSUM62 substitution matrix. This average delta score is the final PROVEAN score^19^. The PROVEAN approach assumes that a variation, that reduces the similarity of a protein to its homologous protein can cause a damaging effect. Thus, the impact of a variation or substitution, on the protein function can be measured as a change in the alignment score that is termed as the delta score for PROVEAN. The low delta scores lead to an increase in the possibility of a deleterious effect on the protein function while the high delta scores are interpreted as variations with neutral effect^32^.

### PolyPhen-2

PolyPhen-2^22,33^ tool coalesce the information of the query sequence, its multiple alignments with homologous proteins and structural parameters to predict the effect of the SNP on the protein function. PolyPhen-2 scans the query sequence in the UniProtKB/Swiss-Prot database using the feature table of the corresponding entry and checks if a given SNP occurs at the functional relevant site i.e. within transmembrane or signal peptide or binding region or not. Similar to SIFT tool, PolyPhen-2 also evaluates the degree of conversion of each substitution position by performing multiple sequence alignment with the homologous sequences. PolyPhen-2 calculates a position-specific independent count (PSIC) score. PolyPhen-2 assumes that a difference of PSIC score between the two variants determines the impact of substitution. Higher PSIC score difference leads to the higher functional impact of the substitution. PolyPhen-2 also performs a BLAST query within the protein structure databases to consider 3D protein structure. Once the 3D structure is found, it is used to investigate the impact of SNP on the structure of the protein (destruction of the hydrophobic core, interaction with legends etc). Eventually, PolyPhen-2 combines all the parameters to apply prediction rules to make the final decision to find whether the SNP has a damaging or benign effect. The SIFT, PROVEAN and PolyPhen-2 tools are available online at http://sift.jcvi.org/, http://provean.jcvi.org/, and http://genetics.bwh.harvard.edu/pph2/dbsearch.shtml respectively.

## RESULTS

### SNP Mining

A total of 2205 records CA 27.29 associated SNPs were found at NCBI SNP database. Among these SNPs, 7 at 3’ splice site, 6 at 5’splice site, 133 at 3’ UTR, 26 at 5’ UTR, and 1351 SNPs in introns were observed. Moreover, 420 SNPs were observed to be missense relative to 13 nonsense SNPs. Consequently, these 2205 SNPs were submitted to dbSNP dataset that categorized 213 (9.66%) SNPs as sSNPs and 24 (1.09%) SNPs as nsSNPs (Fig. 1). These 24 nsSNPs were selected for further investigation. The genomic coordinates and variants of thus filtered 24 nsSNPs were submitted at PROVEAN portal to ‘Human Genome Variants’ tool that provided PROVEAN and SIFT predictions. The default threshold of delta score < = −2.5 was selected to detect the deleterious variations. The list of genomic coordinates and variants of 24 nsSNPs was also submitted at Polyphen 2 portal, by using the option ‘Batch query’. The selected 24 nsSNPs with their nucleotide and amino acid substitutions have been enlisted in table 1. Besides this, overall predictions made by SIFT, PROVEAN, and PloyPhen tools, have been provided for the analyzed 24 nsSNPs. The detail of each variation in nsSNPs was considered to filter the nsSNPs that have a deleterious effect on the protein.

**Table 1.**
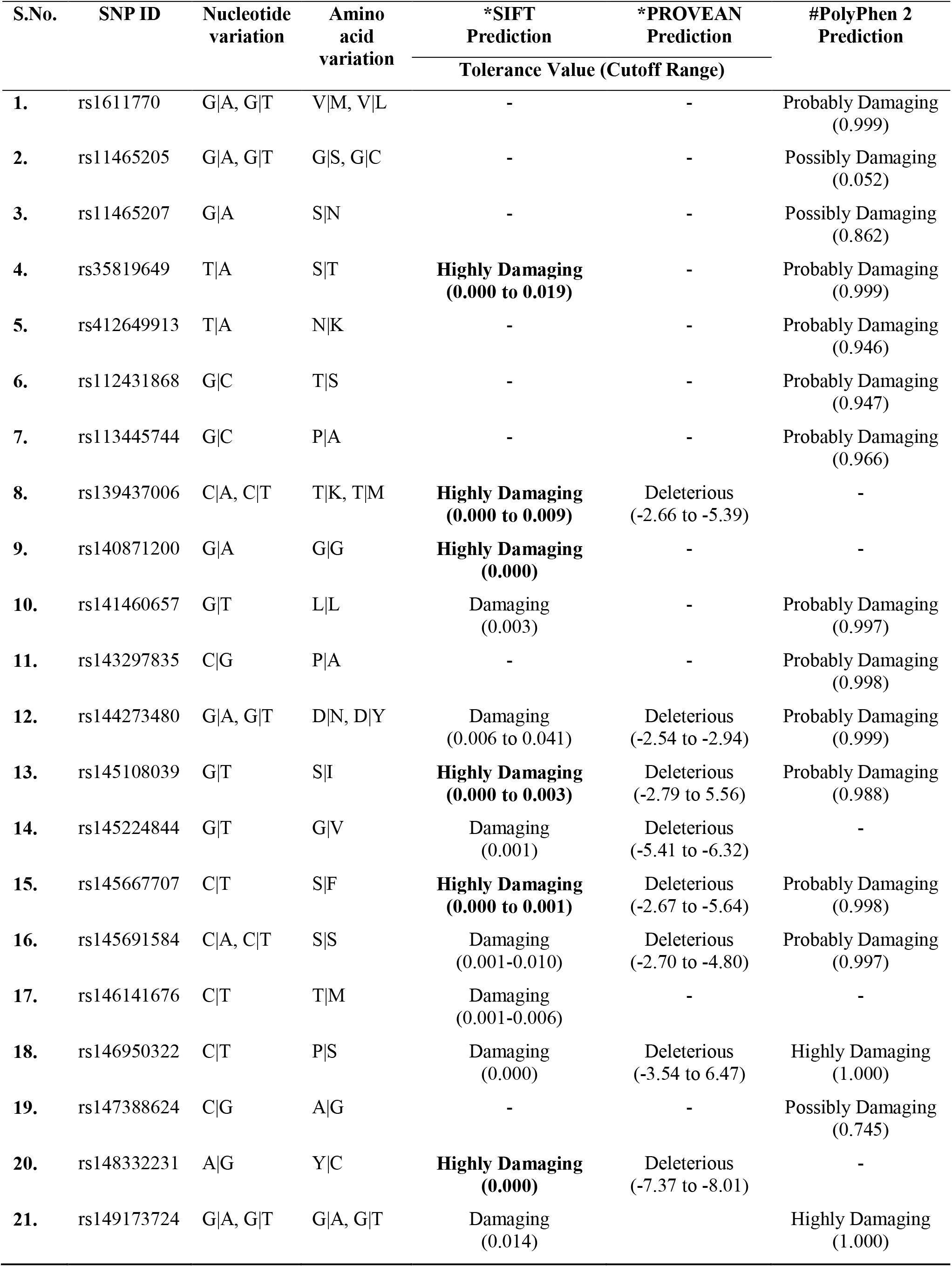
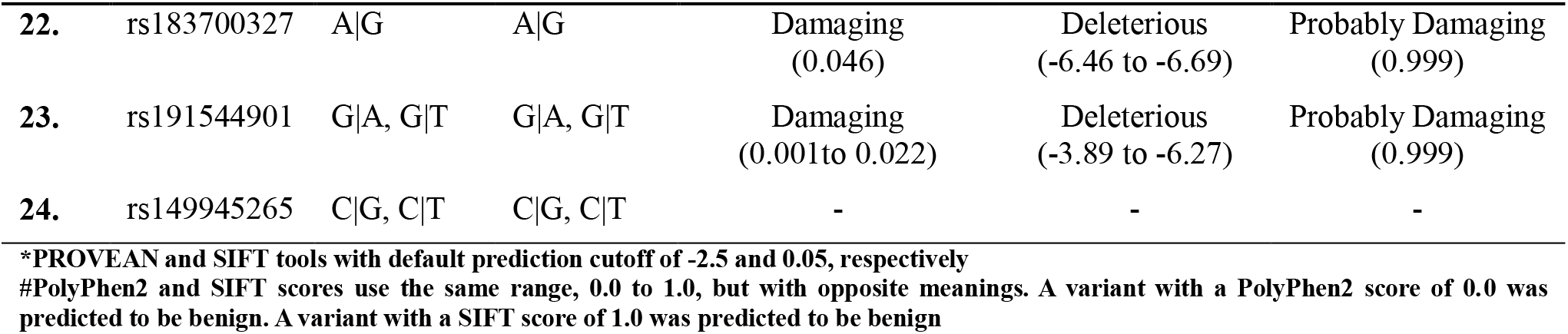
Nucleotide and Amino acid (hydrophobic to hydrophobic) variations in nsSNPs predicted by SIFT, PROVEAN and PloyPhen2 tools

| S.No. | SNP ID | Nucleotide variation | Amino acid variation | *SIFT Prediction | *PROVEAN Prediction | #PolyPhen 2 Prediction |
| --- | --- | --- | --- | --- | --- | --- |
|  |  |  |  | Tolerance Value (Cutoff Range) |  |  |
| 1. | rs1611770 | G\|A, G\|T | V\|M, V\|L | - | - | Probably Damaging (0.999) |
| 2. | rs11465205 | G\|A, G\|T | G\|S, G\|C | - | - | Possibly Damaging (0.052) |
| 3. | rs11465207 | G\|A | S\|N | - | - | Possibly Damaging (0.862) |
| 4. | rs35819649 | T\|A | S\|T | **Highly Damaging (0.000 to 0.019)** | - | Probably Damaging (0.999) |
| 5. | rs412649913 | T\|A | N\|K | - | - | Probably Damaging (0.946) |
| 6. | rs112431868 | G\|C | T\|S | - | - | Probably Damaging (0.947) |
| 7. | rs113445744 | G\|C | P\|A | - | - | Probably Damaging (0.966) |
| 8. | rs139437006 | C\|A, C\|T | T\|K, T\|M | **Highly Damaging (0.000 to 0.009)** | Deleterious (-2.66 to -5.39) | - |
| 9. | rs140871200 | G\|A | G\|G | **Highly Damaging (0.000)** | - | - |
| 10. | rs141460657 | G\|T | L\|L | Damaging (0.003) | - | Probably Damaging (0.997) |
| 11. | rs143297835 | C\|G | P\|A | - | - | Probably Damaging (0.998) |
| 12. | rs144273480 | G\|A, G\|T | D\|N, D\|Y | Damaging (0.006 to 0.041) | Deleterious (-2.54 to -2.94) | Probably Damaging (0.999) |
| 13. | rs145108039 | G\|T | S\|I | **Highly Damaging (0.000 to 0.003)** | Deleterious (-2.79 to 5.56) | Probably Damaging (0.988) |
| 14. | rs145224844 | G\|T | G\|V | Damaging (0.001) | Deleterious (-5.41 to -6.32) | - |
| 15. | rs145667707 | C\|T | S\|F | **Highly Damaging (0.000 to 0.001)** | Deleterious (-2.67 to -5.64) | Probably Damaging (0.998) |
| 16. | rs145691584 | C\|A, C\|T | S\|S | Damaging (0.001-0.010) | Deleterious (-2.70 to -4.80) | Probably Damaging (0.997) |
| 17. | rs146141676 | C\|T | T\|M | Damaging (0.001-0.006) | - | - |
| 18. | rs146950322 | C\|T | P\|S | Damaging (0.000) | Deleterious (-3.54 to 6.47) | Highly Damaging (1.000) |
| 19. | rs147388624 | C\|G | A\|G | - | - | Possibly Damaging (0.745) |
| 20. | rs148332231 | A\|G | Y\|C | **Highly Damaging (0.000)** | Deleterious (-7.37 to -8.01) | - |
| 21. | rs149173724 | G\|A, G\|T | G\|A, G\|T | Damaging (0.014) |  | Highly Damaging (1.000) |
| **22.** | rs183700327 | A\|G | A\|G | Damaging<br>(0.046) | Deleterious<br>(-6.46 to -6.69) | Probably Damaging<br>(0.999) |
| **23.** | rs191544901 | G\|A, G\|T | G\|A, G\|T | Damaging<br>(0.001 to 0.022) | Deleterious<br>(-3.89 to -6.27) | Probably Damaging<br>(0.999) |
| **24.** | rs149945265 | C\|G, C\|T | C\|G, C\|T | - | - | - |

**\*PROVEAN and SIFT tools with default prediction cutoff of -2.5 and 0.05, respectively**
**#PolyPhen2 and SIFT scores use the same range, 0.0 to 1.0, but with opposite meanings. A variant with a PolyPhen2 score of 0.0 was predicted to be benign. A variant with a SIFT score of 1.0 was predicted to be benign**

**Figure 1.**
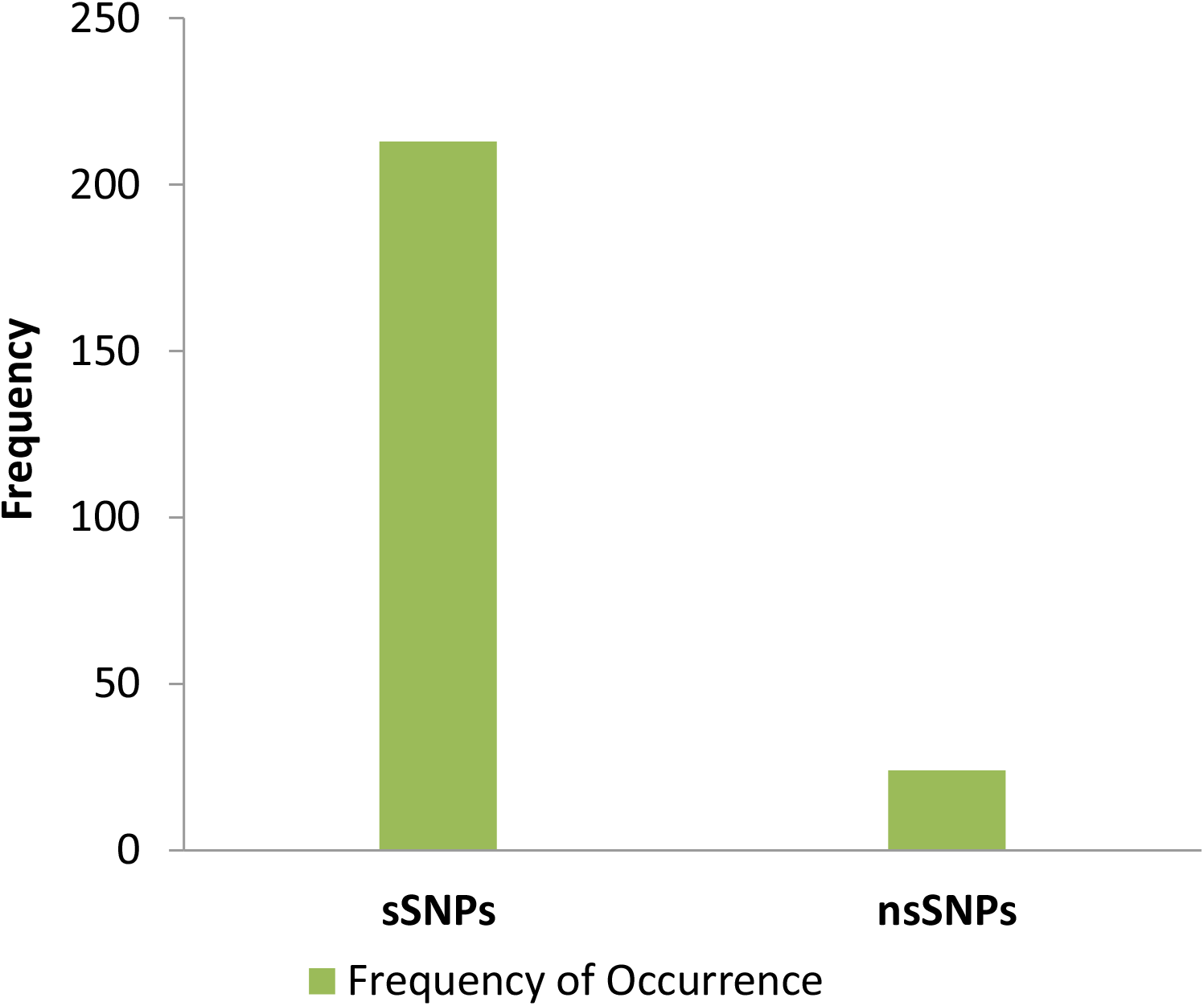
The frequency of occurrence of various sSNPs and nsSNPs associated with CA27.29 gene

### Damaging nsSNPs predicted by SIFT tool

Among 24 analyzed nsSNPs, 9 nsSNPs rs141460657, rs144273480, rs144273480, rs145691584, rs146141676, rs146950322, rs149173724, rs183700327 and rs191544901 were predicted by SIFT tool to be damaging to CA27.29 with a tolerance index score > =0.5 (Table 1). While, seven nsSNPS namely rs35819649, rs139437006, rs140871200, rs145108039, rs145667707, rs148332231 and rs146950322 were predicted to be highly damaging to CA27.29 with a tolerance index score of 0.00 (Table 1). The detailed analysis of each damaging allelic variation in the tested nsSNPs, predicted as functionally significant by SIFT tool has been also provided in Appendix A.

### Deleterious nsSNPs predicted by PROVEAN tool

While SIFT and PROVEAN tools predicted rs139437006, rs145108039, rs145244844, rs145691584, rs146950322, rs148332231, rs183700327, rs191544901 nsSNPs to have adverse effects on the protein (Fig. 2). On the other hand, SIFT and POLYPHEN2 tools predicted rs1611770, rs145667707, rs35819649, rs141460657, rs145224844, rs145691584, rs148332231, rs183700327, rs191544901, rs141460657, rs144273480, rs146950322 nsSNPs to have negative effects on CA27.29 (Fig. 2).

**Figure 2.**
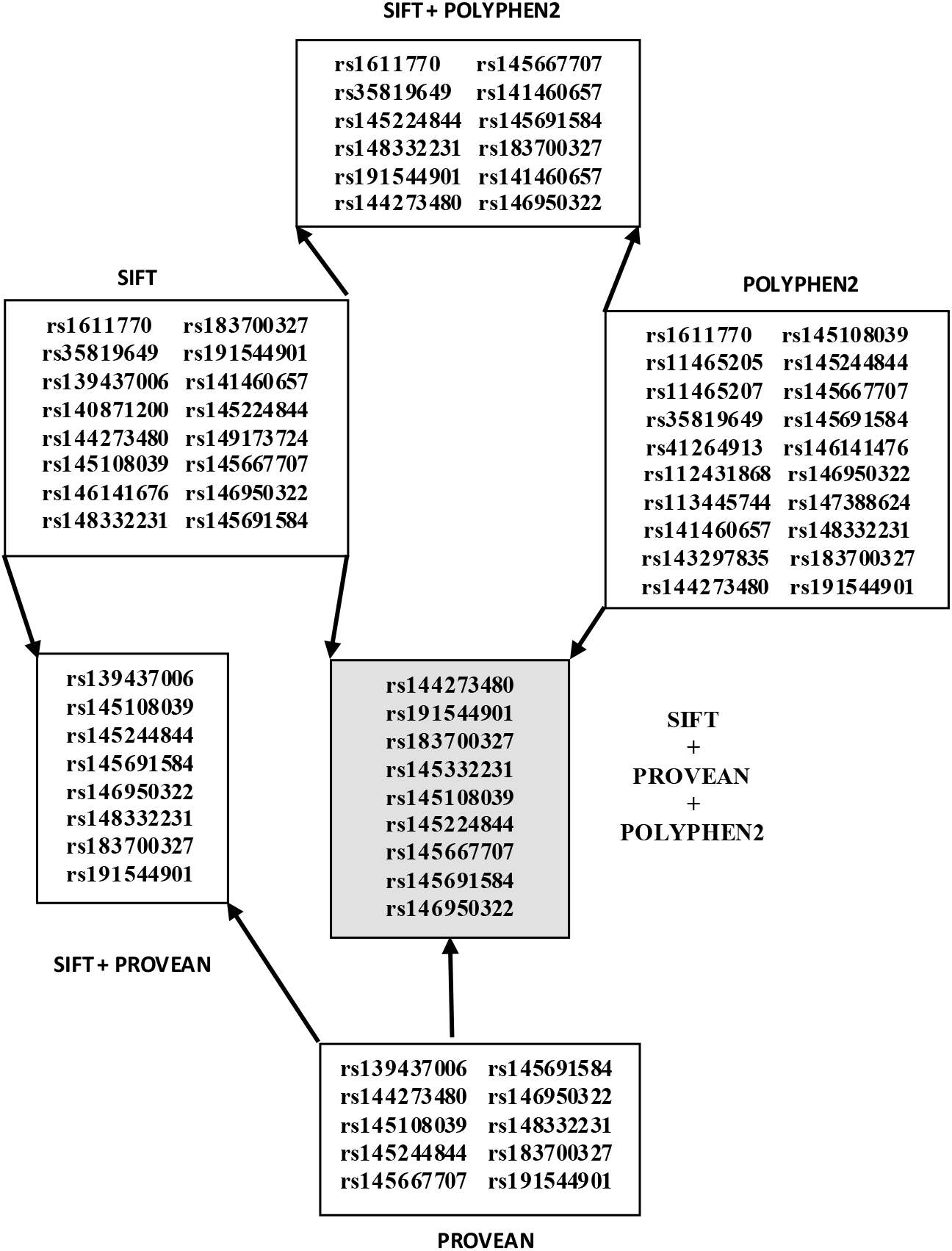
Details of overall nsSNPs predicted by SIFT, PROVEAN and PolyPhen-2 tools

PROVEAN tool predicted 10 nsSNPs rs148332231, rs183700327, rs146950322, rs191544901, rs145224844, rs145667707, rs145108039, rs139437006 and rs145691584 to be highly deleterious to CA27.29 with a delta score of < − 4.00 (Table 1). The detailed analysis of each damaging allelic variation in all the analyzed nsSNPs, predicted as functionally significant by PROVEAN tool has been also provided in Appendix B.

### Damaging nsSNPs predicted by PolyPhen2 tool

Out of 24 nsSNPs submitted to PolyPhen-2 server, 18 nsSNPs were predicted to be damaging wherein 15 nsSNPs were observed to be probably damaging and 3 nsSNPs were possibly damaging. The remaining 6 nsSNPs were predicted to be benign (Table 1).

### Deleterious/Damaging nsSNPs predicted by at least two or three tools

As summarized in Fig. 2, nsSNPs rs144273480, rs191544901, rs183700327, rs148332231, rs145108039, rs145667707, rs145691584 and rs146950322 were predicted to have negative effect on the CA27.29 protein by all the three tools used in the present work (Table 1).

### Functional consequences of targeted nsSNPs

Functional Consequences of all 24 nsSNPs were identified and out of 24, 23 nsSNPs (95.83%) were found to be missense, wherein 21 nsSNPs (87.5%) and 3 nsSNPs (12.5%) were observed to be located in the intronic region, and 3’UTR region respectively.

### Frequency distribution of allelic variations in nsSNPs

An attempt was also made in the present research to find the frequency distribution of allelic variations in the analyzed 24 nsSNPs by using SIFT, PROVEAN and PolyPhen-2 prediction tools. It was observed that G|T substitution was the most common type of allelic variation present in the analyzed 24 nsSNPs. A G|T substitution was observed in 26.67% nsSNPs whereas 10%, 20%, 23.33%, 6.67%, 6.67%, 6.67% nsSNPS were observed with C|A, C|T, G|A, A|G, T|A and C|G substitutions respectively (Fig. 3). The nsSNPs that were predicted by SIFT as damaging exhibited 10.53%, 21.05%, 15.79%, 36.84%, 10.53%, 5.26% allelic variations for C|A, C|T, G|A, G|T, A|G, T|A and C|G substitutions respectively. SIFT tool did not predict any nsSNP with C|G substitution as damaging. While deleterious nsSNPs, as predicted by PROVEAN tool were observed to have 11.11%, 16.67%, 5.56%, 2.22%, 11.11% allelic variations for C|A, C|T, G|A, G|T and A|G substitutions respectively. PROVEAN did not predict any nsSNP with T|A, and C|G substitutions as damaging (Fig. 3).

**Figure 3.**
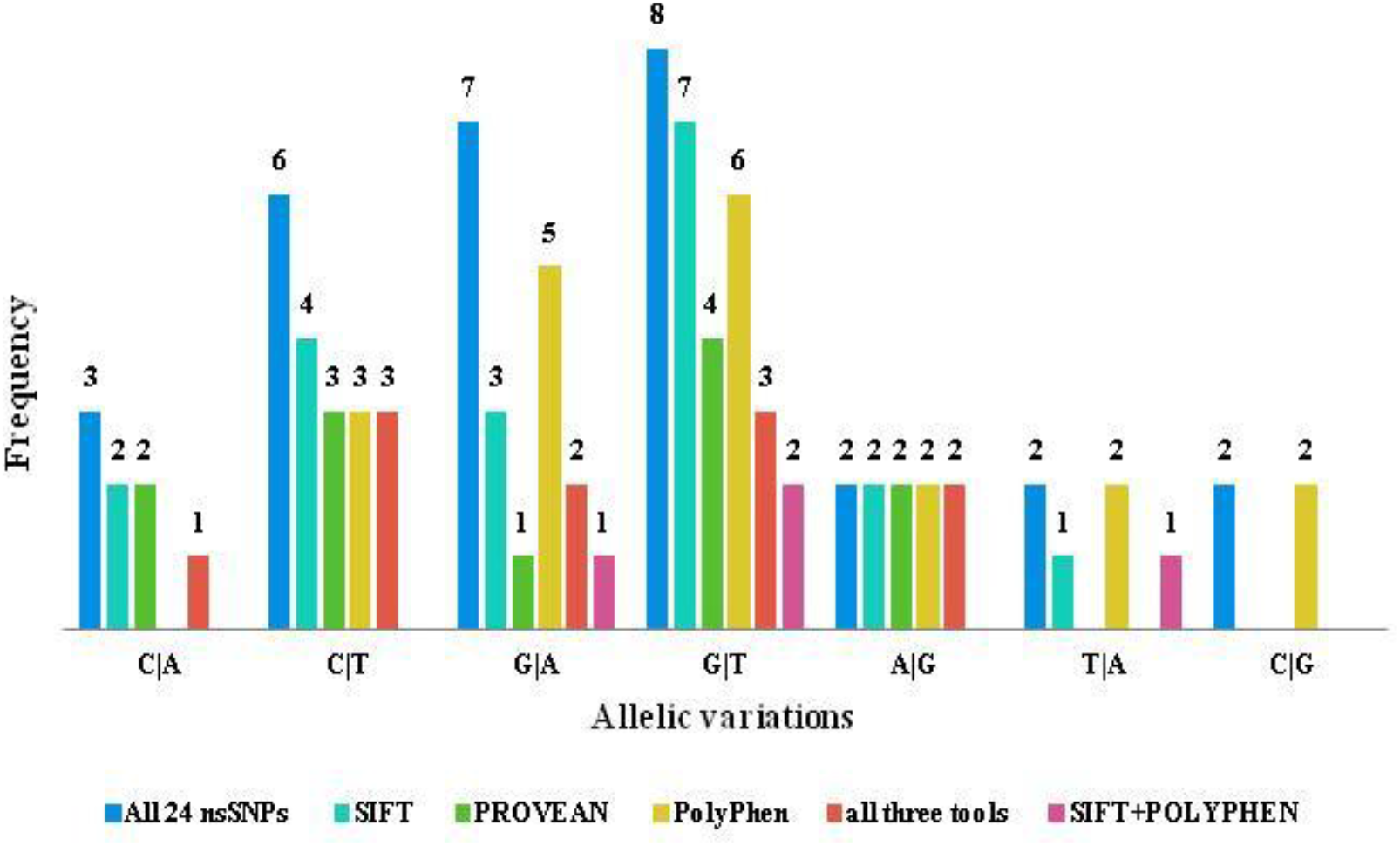
Frequency distribution of allelic variations in deleterious nsSNPs

### Frequency distribution of amino acid to amino acid variations in nsSNPs

It was observed that the analyzed nsSNPs resulting in 100% damage to CA27.29 had one hydrophobic amino acid to another hydrophobic amino acid variation. The details of the amino acid to amino acid variation (AA change) in all the selected nsSNPs have been provided in Table 1. The status of the polarity of amino acid variations was also analyzed (Fig. 4). It was found that the polarity of amino acid in 15 (62.5%) nsSNPs was not affected even if the observed variation was damaging. Wherein 8 (33.33%) and 7 (29.17%) nsSNPs were analyzed with the polar amino acid to polar amino acid (P to P) and non-polar amino acid to non-polar amino acid (NP to NP) variations respectively (Fig. 4). However, non-polar to polar (NP to P) and polar to non-polar (P to NP) amino acid variations were observed to occur in 6 (25%) and 3 (12.5%) nsSNPs respectively. The variations of polarity distribution in nsSNPs, predicted by SIFT tool to be damaging, suggested that NP to P amino acid variations had the maximum frequency of 33.33% followed by NP to NP (27.78%), P to P (27.78%) and P to NP (11.11%) amino acid variations. While polarity distribution of variations in nsSNPs, predicted deleterious by PROVEAN tool, suggested that the frequency (36.36%) of P to P amino acid variations was observed to be maximum. However, the frequency of NP to P and P to NP amino acid variations in deleterious nsSNPs were assessed to be 27.27% and 9.09% respectively. The damaging variation as predicted by PolyPhen2 exhibited the frequency of 31.58% for NP to NP and P to P amino acid variations. However, other NP to P and P to NP variations in polarity distributions were noted to have the frequency of 10.53% and 26.32% respectively (Fig. 4).

**Figure 4.**
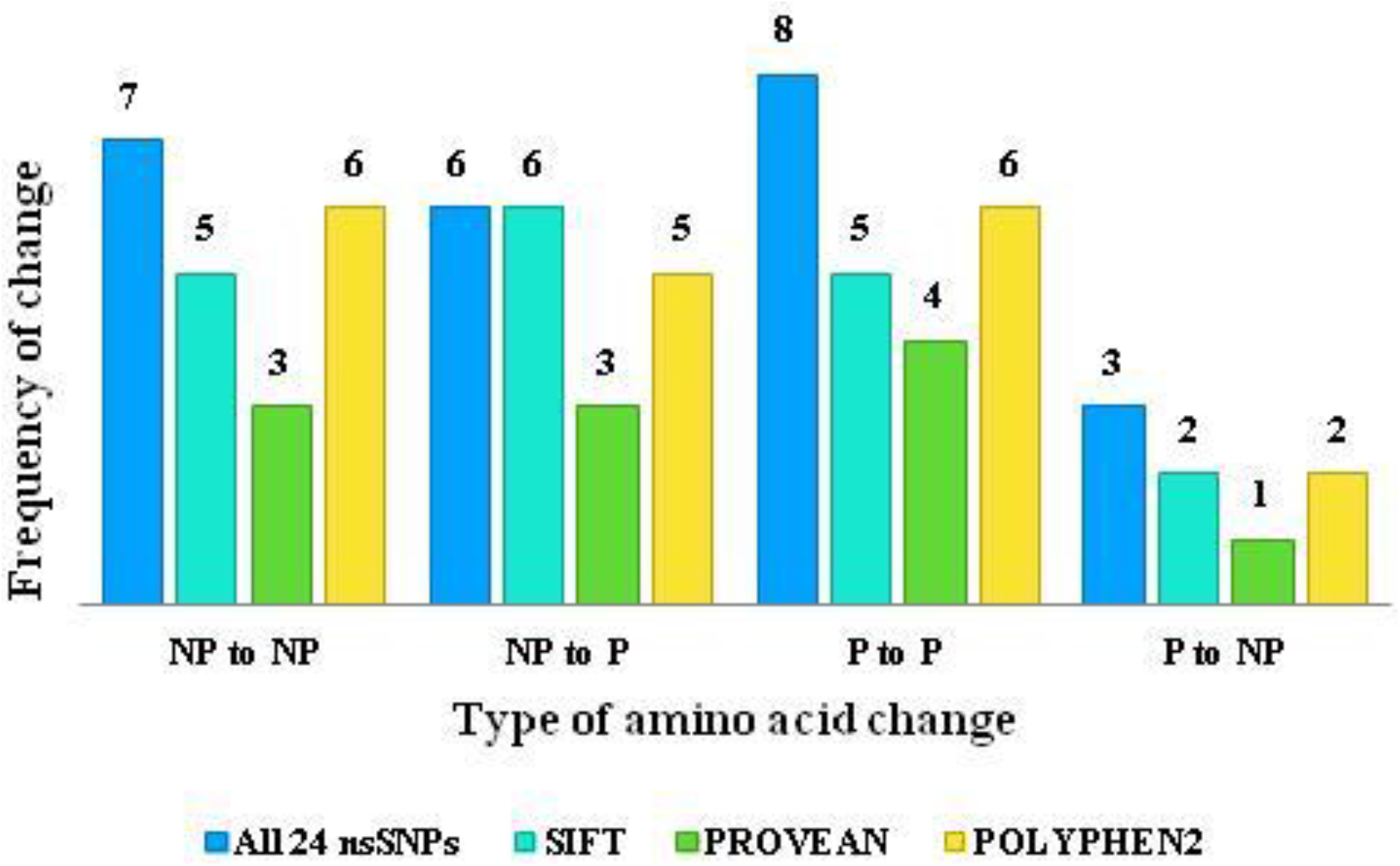
Frequency distribution of amino acid to amino acid (NP= Non-Polar; P= Polar) variations in deleterious nsSNPs

### Benchmarking of the results provided by SIFT and PROVEAN tools using clustering methods

Four clustering methods KMeans^34^, MakeDensityBasedClustrer^35^, FarthestFirst^36,37^ and ACERAM^38^ were used to benchmark the number of damaging or deleterious nsSNPs identified by SIFT, POLYPHEN2, and PROVEAN tools. The method k-means clustered 80% nsSNPs and 66% nsSNPs correctly, that were predicted as damaging by SIFT and PROVEAN tools respectively (fig.5). While Make_Density_Based_Clustrer clustered 98% and 73% nsSNPs correctly that were predicted as damaging or deleterious by SIFT and PROVEAN tools respectively. On the other hand, FarthestFirst clustered 99% and 83% nsSNPs correctly for the predictions made by SIFT and PROVEAN tools respectively (Fig. 5). As compared to that, ACERAM clustered 99% and 98% nsSNPs correctly, predicted as deleterious by SIFT and PROVEAN tools respectively. Overall results from this experiment indicated that the clustering method ACERAM provided more precise results than the other clusterers used in the present study (Fig. 5).

**Figure 5.**
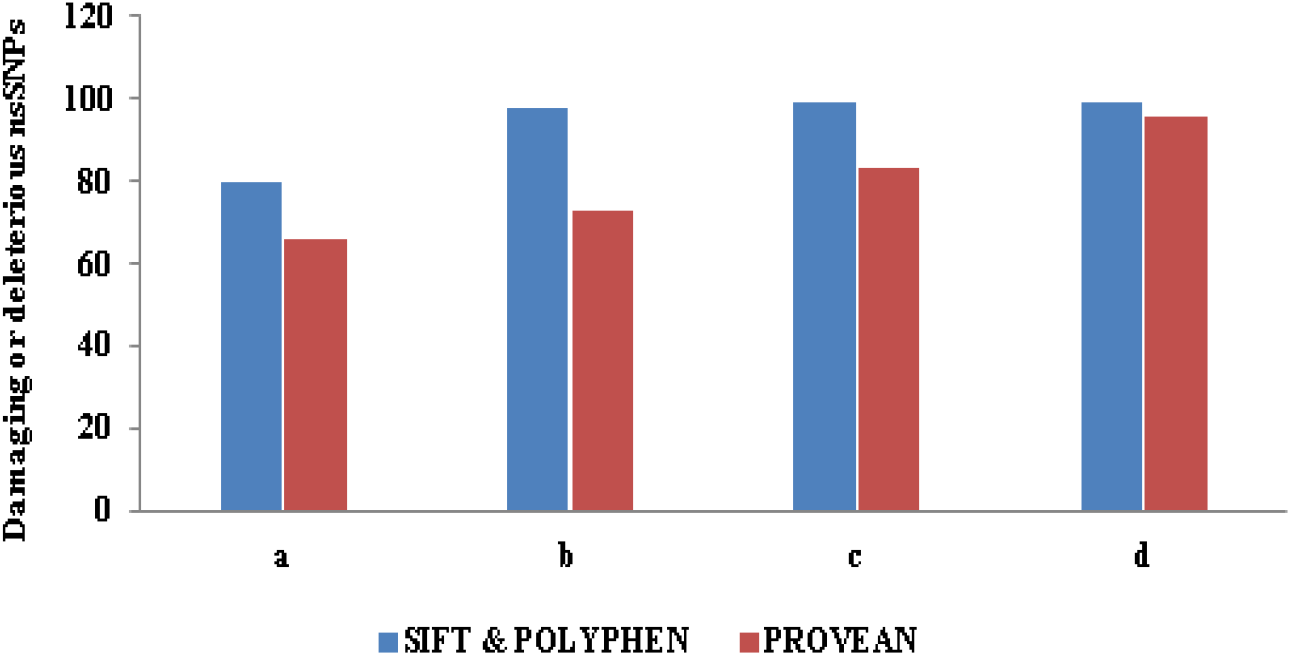
Benchmarking of predictions made by SIFT, PolyPhen2 and PROVEAN tools using clustering methods a. K_Means b. Make_Density_Based_Clustrer c. Farthest_First d. ACERAM

### Comparison of the predictions made by SIFT and PROVEAN tools with SNPs database

The predictions made by computational tools were compared using four clustering methods with SNP database (Table 2). The overall comparisons identified 215 SNPs as damaging whereas 147 SNPs were found to be tolerated by SIFT tool. While the PROVEAN tool predicted 199 SNPs as neutral and 163 SNPs as deleterious. While clustering method K-means revealed that there were 172 damaging, 189 tolerated and 80 SNPs clustered incorrectly (false positive + false negative) predicted by SIFT tool. Similarly, the K-means clustering method identified 168 SNPs as neutral and 193 SNPs as deleterious for the predictions made by the PROVEAN tool, wherein, 66 SNPs were recognized as incorrectly clustered (Table 2).

ACERAM clusterer identified 201 and 122 SNPs as damaging and tolerated respectively. A total of 69 SNPs were marked as incorrectly clustered for the predictions made by the SIFT tool. While for the predictions made by the PROVEAN tool, ACERAM identified 167 SNPs as neutral and 156 as deleterious SNPs. ACERAM clusterer observed, 69 SNPs as incorrectly clustered for the predictions made by the PROVEAN tool (Table 2). On the other side, Make_Density_Based_Clusterer method clustered 274 SNPs as damaging and 87 SNPs as tolerated whereas 98 SNPs were found to be incorrectly clustered as predicted by SIFT tool. The predictions made by the PROVEAN tool provided 161 SNPs as neutral, 200 deleterious while 73 SNPs as incorrectly clustered. On the other hand, FarthestFirst algorithm clustered 285 SNPs as damaging, 76 tolerated and 99 SNPs as incorrectly clustered for the predictions made by SIFT tools. While the predictions made by PROVEAN tool presented 159 SNPs as neutral and 202 SNPs deleterious and 83 SNPs as incorrectly clustered (Table 2).

**Table 2.** Comparison of the prediction made by SIFT and PROVEAN tools using various clusterers

| Method | SIFT |  |  | PROVEAN |  |  |
| --- | --- | --- | --- | --- | --- | --- |
|  | Damaging | Tolerated | Incorrectl<br>y<br>Clustered | Neutral | Deleterious | Incorrectly<br>Clustered |
| SNPs |  |  |  |  |  |  |
| K-Means | 172 | 189 | 80 | 168 | 193 | 66 |
| ACERAM | 201 | 122 | 92 | 167 | 156 | 69 |
| Make_Density_<br>Based_Clusterer | 274 | 87 | 98 | 161 | 200 | 73 |
| FarthestFirst | 285 | 76 | 99 | 159 | 202 | 83 |

### Validation of polymorphisms in nsSNPS using SNP database

It is important to check the true status of polymorphism and confirm validation status for the analyzed nsSNPs. A polymorphism can be validated by independent submissions of the frequency or genotypic data to the dbSNP database^19^. The genotypes of each of the 24 nsSNPs were submitted one by one to dbSNP server to observe their validation. The functional consequences for each of the nsSNPs were checked and validated by 1000G clusterer, Heapmap and frequency cluster validation techniques. Global MAF and clinical significance of the analyzed nsSNPs were also observed in the present study. The observations derived from these analyses have been provided in Table 3.

**Table 3.**
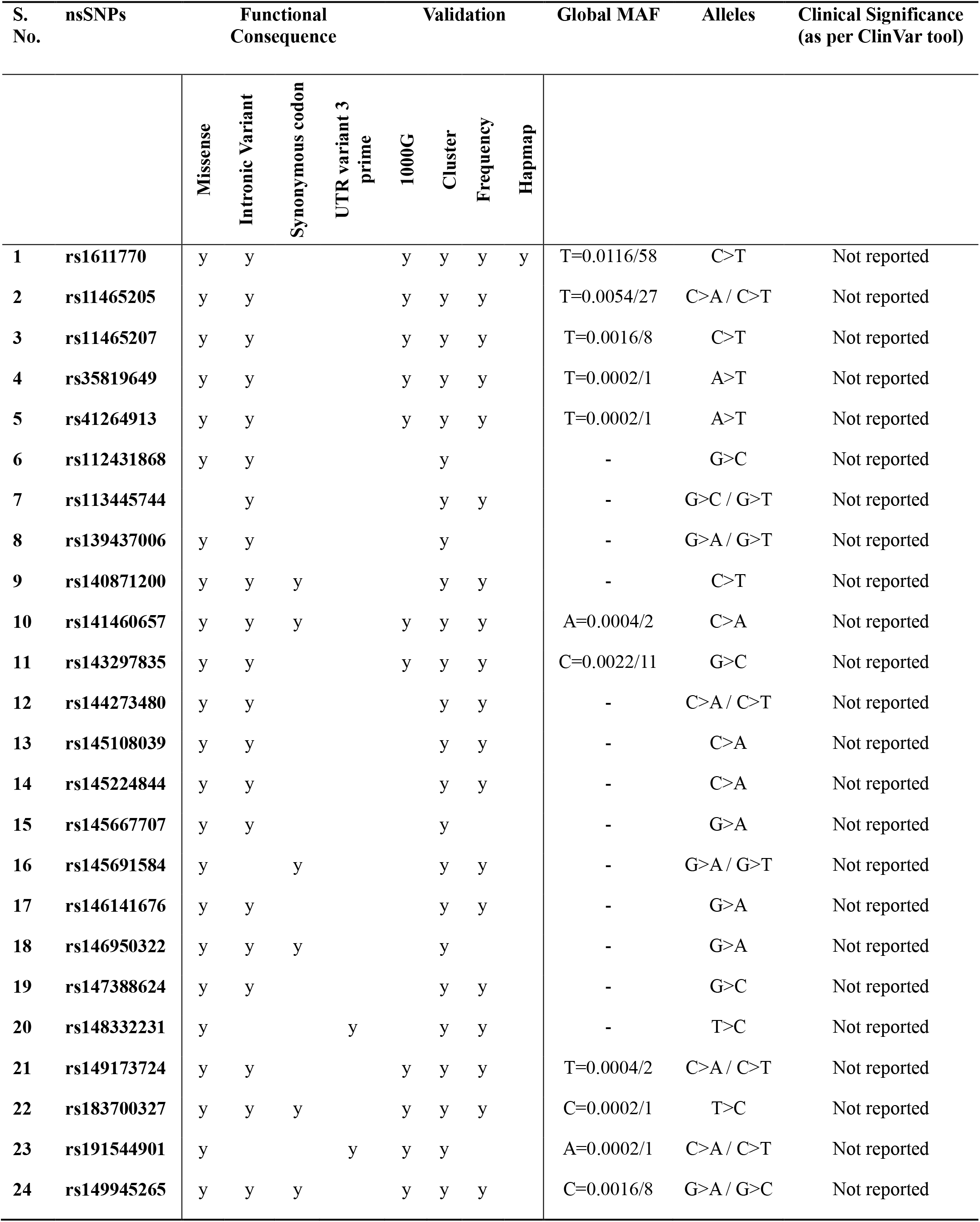
Functional consequences, validation, global MAF and clinical significance of the observed nsSNPs

## DISCUSSION

To distinguish the critical regions of a gene, it is important to be familiar with its genetic variation. SNPs are genetic variations that have been linked and reported with various complex diseases such as sickle-cell anemia, β-thalassemia, cystic fibrosis and breast cancer etc.^19^. For example, SNPs in BRCA-1 gene are not just involved in the onset of the disease but are also involved in promoting the disease progression^39,40^. In the present study, we analyzed the MUC1 gene product CA27.29 protein to identify possible nsSNPs that modify its functional properties. There are various features used by computational tools to predict the functional influences of nsSNPs on the gene product. In the present work, three contemporary online computational tools, SIFT^21,27,28^, Provean^22,30,31^ and PolyPhen2^23,33^ were used, to predict the effects of different allele variations in nsSNPs on amino acid substitution and protein function. Using SIFT, PROVEAN and POLYPHEN2 tools, out of 2205 analyzed nsSNPs, a total of 24 (1.08%) malicious nsSNPs were predicted. These 24 nsSNPs were analyzed further wherein 9 (37.5%) out of 24 nsSNPs were predicted as damaging or deleterious. SIFT tool predicted 16 (66.66%) nsSNPs as damaging while PROVEAN tool predicted 10 (41.66%) nsSNPs as deleterious. Whereas PolyPhen2 tool predicted 20 (83.33%) nsSNPs as damaging. A total of 14 (58.33%) nsSNPs were predicted to have a damaging effect when at least two predictions tools were used. Intriguingly, 12 (50%) nsSNPs were predicted in common to be deleterious by SIFT and PolyPhen2 tools. This might be due to the common steps of assessing the degree of the conversation by both the tools that utilize multiple sequence alignment of the homologous sequences^19^. Similarly, a total of 6 (25%) nsSNPs were predicted in common by SIFT and PROVEAN tools. Ankyrin (ANK) repeat motifs are unique motif mediating protein-protein interaction, the structure of which has been observed to be conserved and known to occur in many functionally assorted proteins. But in the present study, out of 24 analyzed nsSNPs, none of nsSNPs was found to contain the ANK repeat motif.

There are many misclassified SNPs in the public databases, therefore it was important to validate the nsSNPs under present investigation. On submitting the selected 24 nsSNPs under investigation to dbSNP, importantly, the allele frequencies were provided for only 11 nsSNPs. SNPs can be classified according to their minor allele frequencies wherein the variants with minor allele frequencies (MAF) between 0.1% and 3% are defined as rare variants while the variants with minor allele frequencies of less than 0.1% are defined as novel variants. While the variants with high allele frequencies greater than 5% are defined as common variants^25,34^. It is important to note that the minor allele frequency of all the analyzed nsSNPs in the present study was observed to be below 0.1% which establish these variants as novel variants that are yet to be studied. The identification of the functional consequences of 24 nsSNPs revealed that out of 24 analyzed nsSNPs, 23 (95.83%) nsSNPs are missense. It has been reported that the cancer-associated missense mutations can lead to the drastic destabilization of the resulting protein^41^. Besides this, it was also assessed in the present study, that 21 (87.5%) nsSNPs were located in the intronic region and 3 (12.5%) nsSNPs were found to be located in 3’UTR region. The introns are removed in the process known as RNA splicing in mature RNA, that suggests that these will not be expressed in the final messenger RNA (mRNA) product. Thus 21 (87.5%) nsSNPs located in the intronic region might be having no functional consequence on CA27.29 protein but probably the rest of 3 (12.5%) nsSNPs rs145691584, rs148332231 and rs191544901 might have the functional role. To check the clinical significance of the selected 24 nsSNPs, ClinVar tool was used and the observations showed that all of the analyzed nsSNPs are yet to be reported. While assessing the frequency distribution of allelic variations, G to T allelic variation was found to occur in 8 (33.33%) out of the 24 nsSNPs tested followed by G to A substitution, frequency of which was observed in 7 (29.16%) nsSNPs under investigation. Similarly, SIFT and PROVEAN tools also predicted allelic variation in 7 (36.84%) and 4 (33.33%) nsSNPs with G->T substitutions while C->T substitutions were observed in 4 (21.05%) and 3 (25%) deleterious nsSNPs. Polyphen2 tool predicted allelic variation in 6 (30%) nsSNPs with G->T substitution while G->A substitution was observed in 5 (25%) nsSNPs. In light of these observations, it was assumed that probably G->T/A and C/G->T allelic variations are responsible for the damaging effect of the analyzed nsSNPs. Four different clustering algorithms k-means, Make_Density_Based_Clustrer, Farthest_First and ACERAM were used to compare the predictions made by SIFT and PROVEAN tools for each allelic variation. SIFT tool made incorrect predictions for 80 nsSNPs while it was observed to be 66 with PROVEAN tool when assessed with k-means clustering algorithm. However, using ACERAM clustering algorithm, SIFT and PROVEAN tools were found to made 92 and 69 incorrect predictions respectively. Similar to this, Make_Density_Based_Clusterer identified 98 and 73 incorrect predictions in nsSNPs variations made by SIFT and PROVEAN tools respectively. On the other hand, as per FarthestFirst clustering algorithm, 99 and 83 nsSNPs variations were differently predicted by SIFT and PROVEAN tools respectively. The present analysis suggest that some of the allelic variations were misclassified and the predictions made by these tools are not 100% accurate. Consequently, investigating the prediction accuracy of these tools is an area of further research. Or it could be also possible that there are few incorrect or incomplete sequences submitted in the public databases that have led these tools to made predictions in the wrong direction. To find out these incomplete and incorrect sequences in SNP database is another prospective research area.

## CONCLUSIONS

After primary treatment of breast cancer, the level of MUC1 gene product CA27.29 in the bloodstream can predict the disease recurrence about six months in advance. For this and many other reasons, it was important to know the factors affecting its stability, particularly the functionality of MUC 1 gene and its associated CA27.29 glycoprotein. Besides this, few reports directly relate nsSNPs to the increased risk of breast cancer. To keep these in mind an attempt was made in the present study to identify nsSNPs that have malicious effect on CA27.29 protein. From a total of 2205 SNPs recovered from dbSNP database, 24 nsSNPs were found to be related to the target. SIFT, PROVEAN and Polyphen2 tools predicted 16, 10 and 20 numbers of nsSNPs as deleterious or damaging respectively. Subsequent to this analysis, in the C/G->T and G->A/T allelic variations were prominently observed in the deleterious nsSNPs under investigation. The clinical significance as assessed by ClinVar tool confirmed that all the analyzed nsSNPs are yet to be reported and are not studied till date. The results were validated after submitting nsSNPs individually to the dbSNP to find their Global minor allele frequency (Global MAF) that revealed allele frequencies for only 11 nsSNPs. In addition, the allelic frequency of all nsSNPs was observed to be below 0.1% that might establish to consider these nsSNPs as the novel variants. Moreover, 3 nsSNPs rs145691584, rs148332231 and rs191544901 were found to be located in 3’UTR region of CA27.29 gene that might have the functional role in altering its protein levels in the blood stream of breast cancer patients.

**Appendix A.**
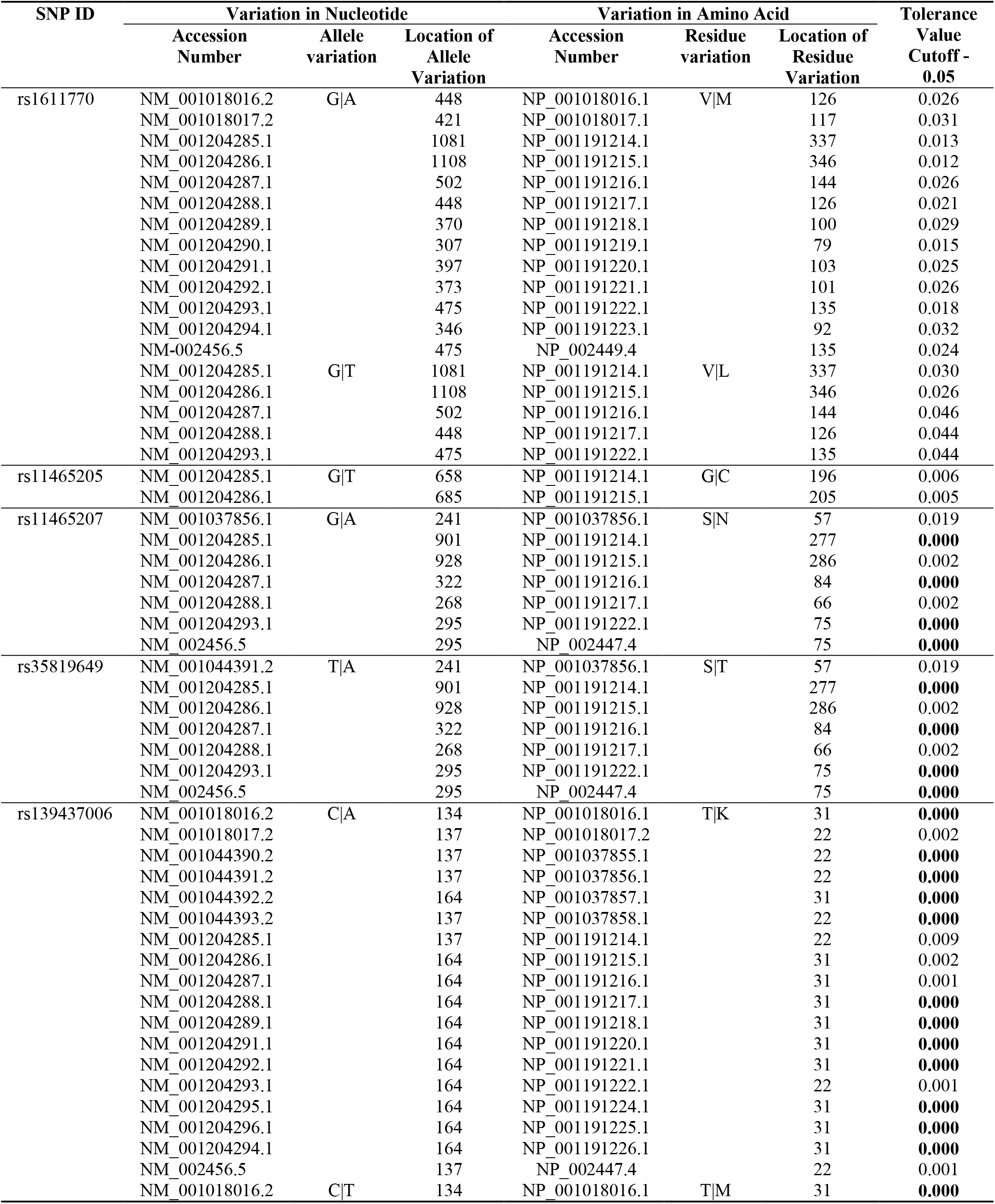
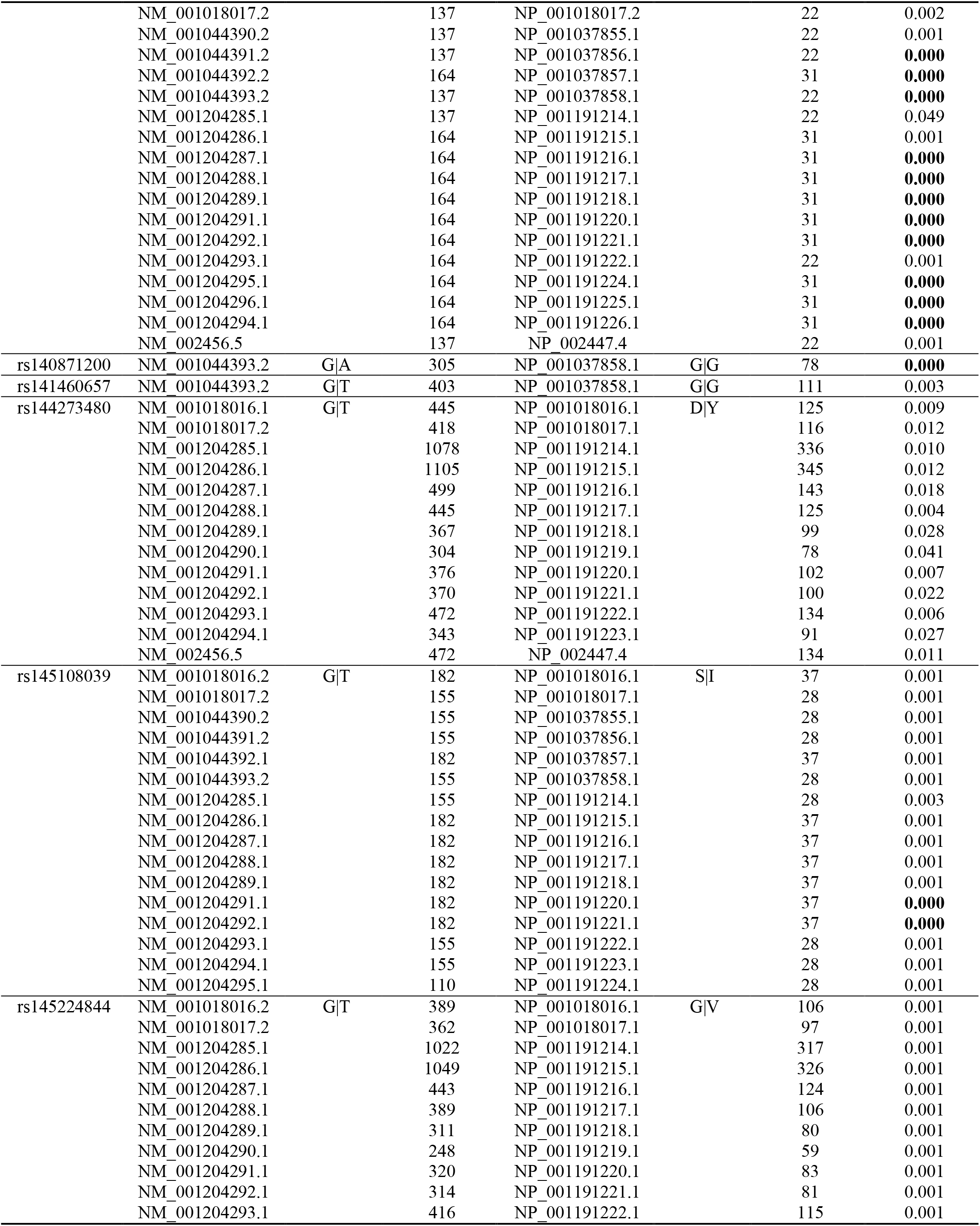
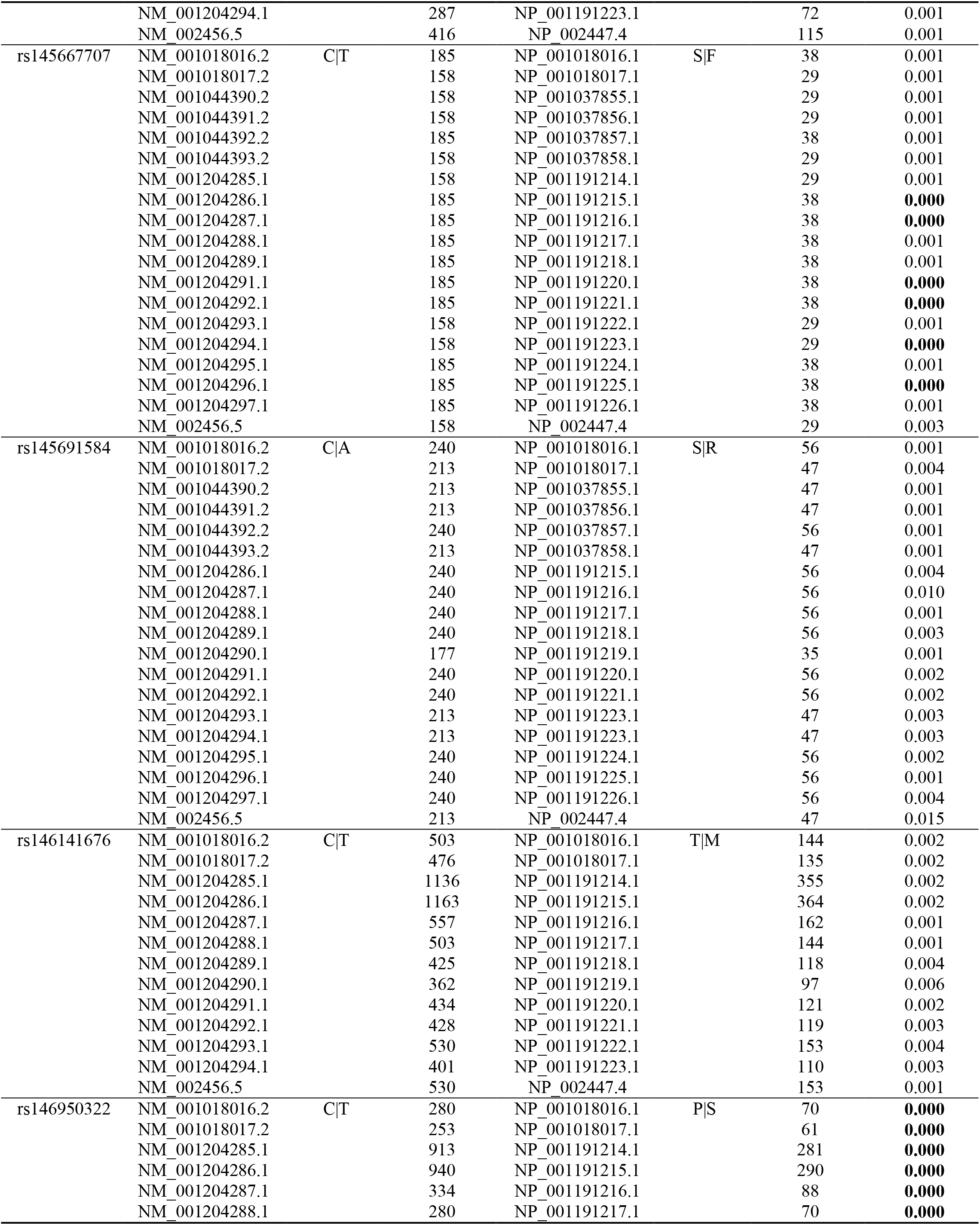
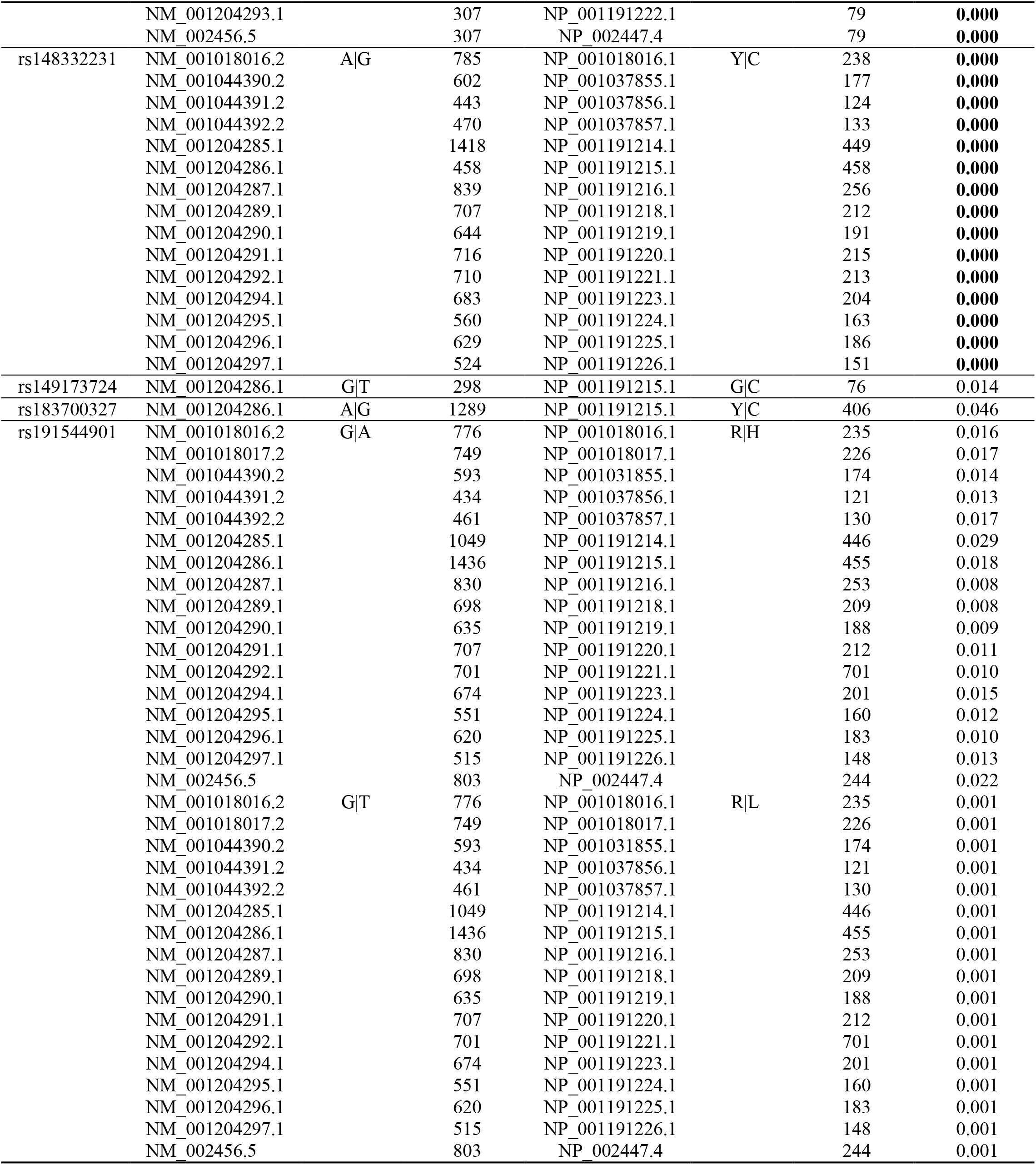
Summary of deleterious nsSNPs predicted by SIFT tool.

**Appendix B.**
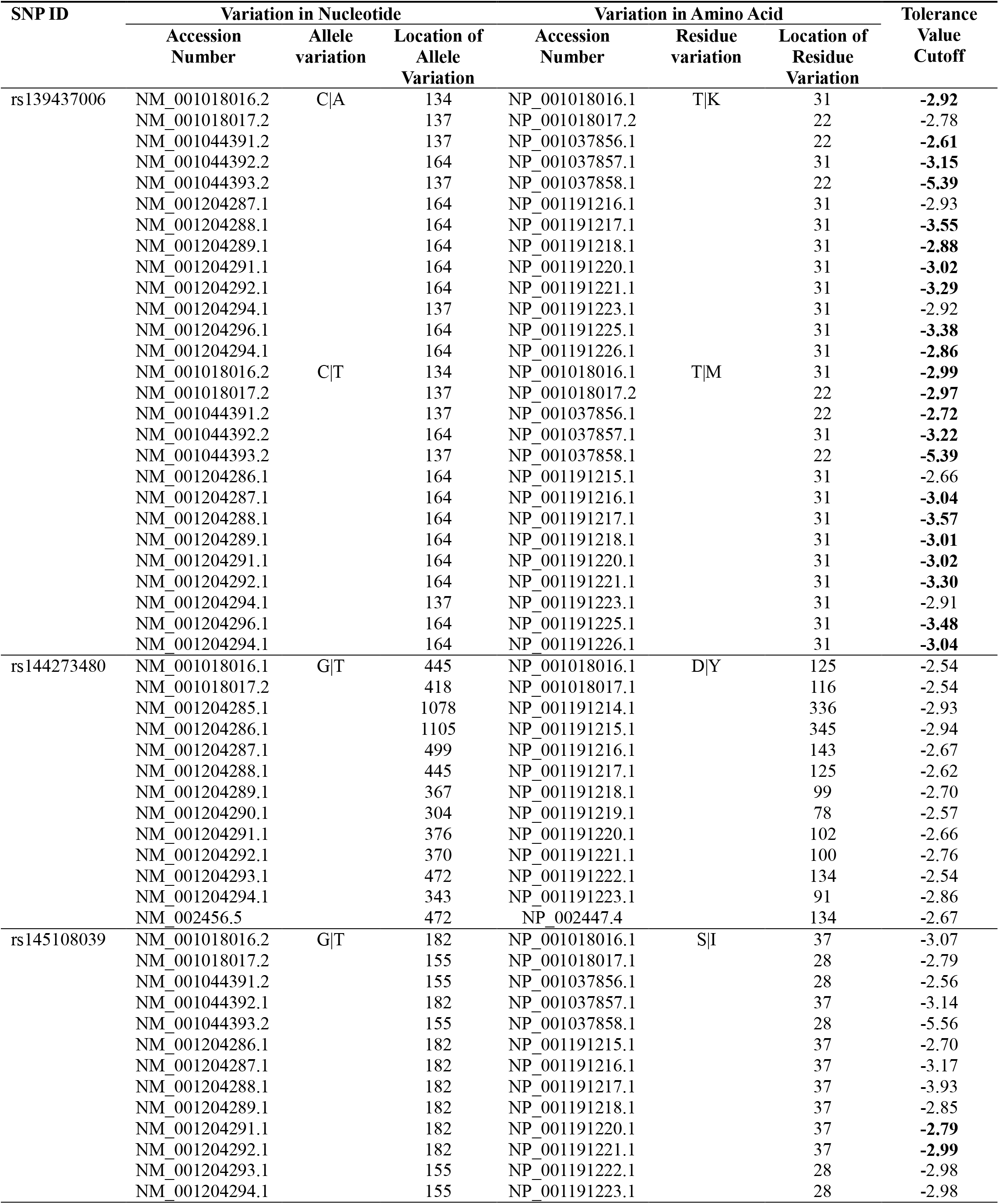
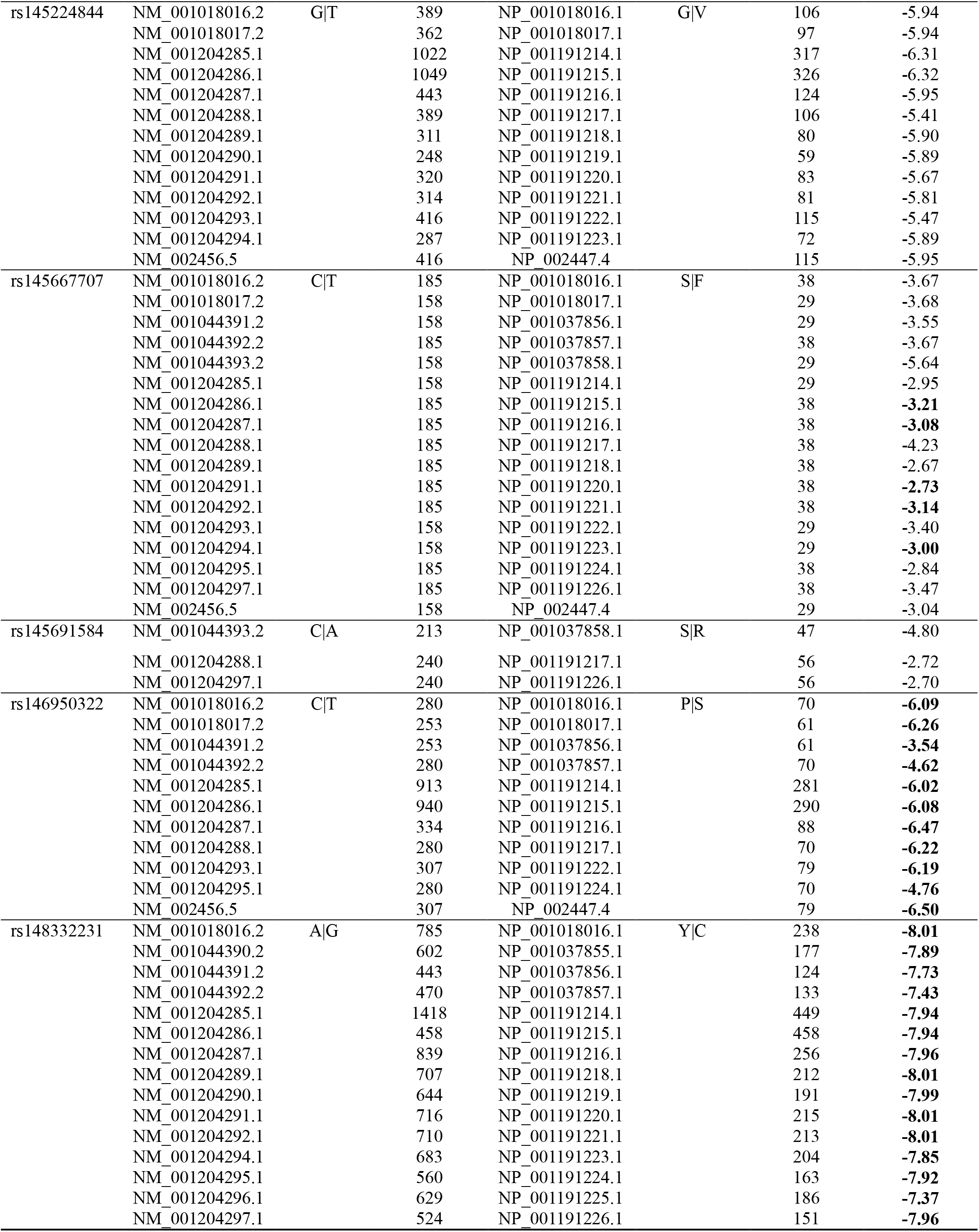
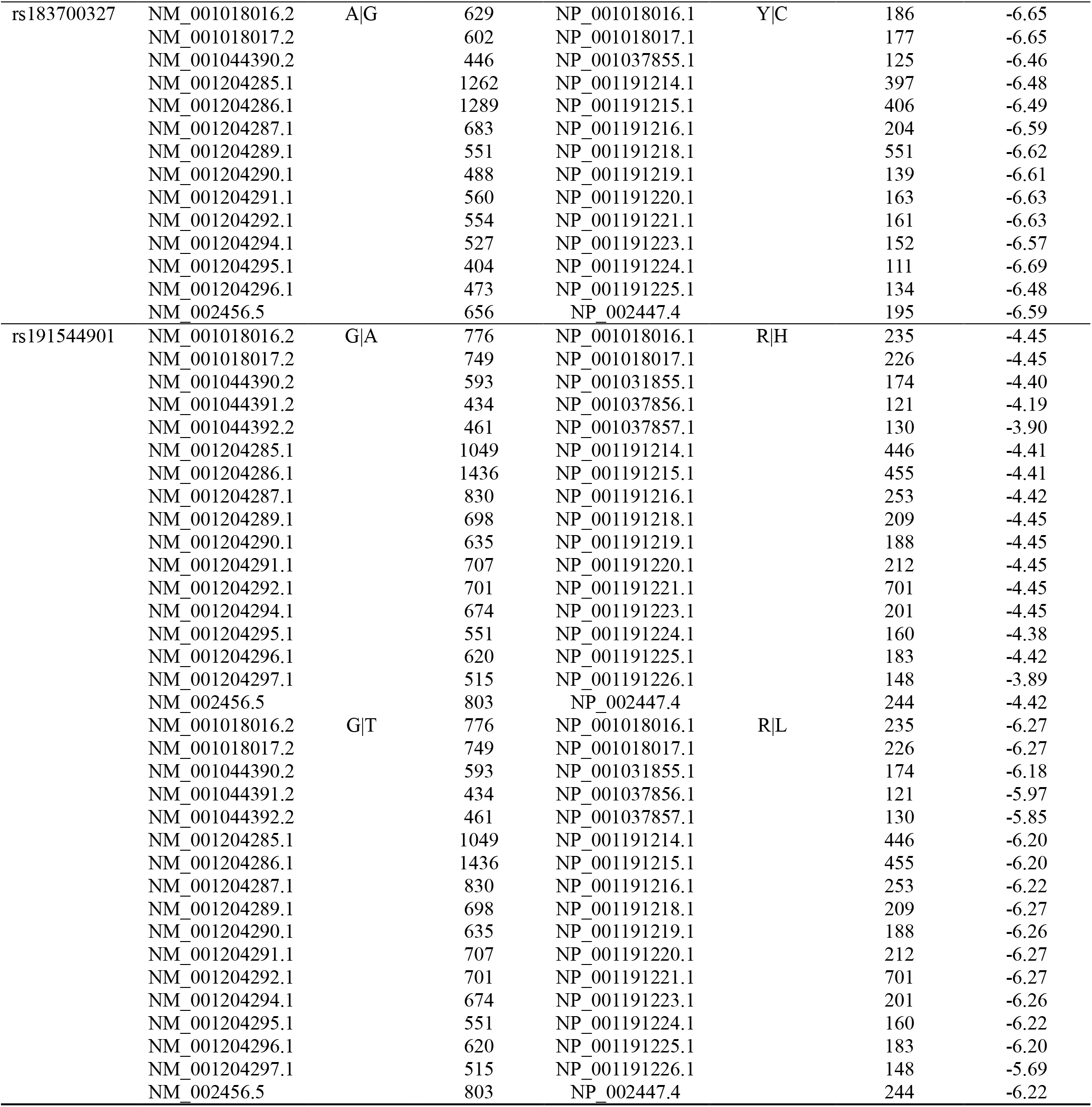
Summary of deleterious nsSNPs predicted by PROVEAN tool.

## REFERENCES

1. Saad A, Abraham J. Role of tumor markers and circulating tumors cells in the management of breast cancer. Oncology 2008; 22(7):726–31.

2. Bray F, Ferlay J, Soerjomataram I, Siegel RL, Torre LA, Jemal A. Global cancer statistics 2018: GLOBOCAN estimates of incidence and mortality worldwide for 36 cancers in 185 countries. CA Cancer J Clin 2018; 68(6):394–424.

3. American Cancer Society. 2017. Breast Cancer Facts & Figures 2017-2018. Atlanta: American Cancer Society, Inc.

4. Price MR, Hudecz F, O’Sullivan C, Baldwin RW, Edwards PM, Tendler SJ. Immunological and structural features of the protein core of human polymorphic epithelial mucin. Mol Immunol 1990; 27(8):795–802.

5. Briggs S, Price MR, Tendler SJ. Immune recognition of linear epitopes in peptide fragments of epithelial mucins. Immunology 1991; 73(4):505–507.

6. Harris L, Fritsche H, Mennel R, Norton L, Ravdin P, Taube S, et al. American Society of Clinical Oncology 2007 update of recommendations for the use of tumor markers in Breast Cancer. J Clin Oncol 2007; 25(33):5287–5312.

7. Duffy MJ. Serum tumor markers in breast cancer: are they of clinical value?. Clin Chem 2006; 52(3):345–351.

8. Beveridge RA. Review of clinical studies of CA 27.29 in breast cancer management. Int J Biol Markers 1999; 14(1): 36–39.

9. Rakha EA, Boyce RW, Abd El-Rehim D, Kurien T, Green AR, Paish EC, et al. Expression of mucins (MUC1, MUC2, MUC3, MUC4, MUC5AC and MUC6) and their prognostic significance in human breast cancer. Mod Pathol 2005; 18(10): 1295–1304.

10. Kokko R, Holli K, Hakama M. CA15-3 in the follow-up of localised breast cancer: a prospective study. Eur J Cancer 2002; 38(9):1189–1193.

11. Nicolini A, Tartarelli G, Carpi A, Metelli MR, Ferrari P, Anselmi L, et al. Intensive postoperative follow-up of breast cancer patients with tumour markers: CEA, TPA or CA15.3 vs. MCA and MCA-CA15.3 vs. CEA-TPA-CA15.3 panel in the early detection of distant metastases. BMC Cancer 2006; 6(1):269.

12. Rack B, Schindlbeck C, Jueckstock J, Genss EM, Hepp P, Lorenz R, et al. Prevalence of CA 27.29 in primary breast cancer patients before the start of systemic treatment. Anticancer Res 2010; 3(1): 1837–1842.

13. Kabel AM. Tumor markers of breast cancer: New prospectives. J Oncol Sci 2017; 3(1):5–11.

14. Barnes MR. Genetic variation analysis for biomedical researchers: a primer. In: Barnes, MR, G Breen, (Ed.). Genetic Variation. Humana, Press, Totowa, NJ: Methods Molecular Biology; 2010. p. 1–20.

15. Kruglyak L, Nickerson DA. Variation is the spice of life. Nat Genet 2008; 27(3):234–236.

16. Xavier RJ, Rioux JD. Genome-wide association studies: a new window into immune-mediated diseases. Nat Rev Immunol 2008; 8(8): 631–643.

17. Bodmer W, Bonilla C. Common and rare variants in multifactorial susceptibility to common diseases. Nat Genet 2008; 40(6):695–701.

18. Wang JB, Pang GSY, Chong SS, Lee CGL. SNP web resources and their potential applications in personalized medicine. Curr Drug Metab 2012; 13(7): 978–990.

19. Kosaloglu Z, Bitzer J, Halama N, Huang Z, Zapatka M, Schneeweiss A, et al. In silico SNP analysis of the breast cancer antigen NY-BR-1. BMC Cancer 2016; 16(1):901.

20. Nakken S, Alseth I, Rognes T. Computational prediction of the effects of non-synonymous ingle nucleotide polymorphisms in human DNA repair genes. Neuroscience 2007; 145(4):1273–1279.

21. Kumar P, Henikoff S, Ng PC. Predicting the effects of coding non-synonymous variants on protein function using the SIFT algorithm. Nat Protoc 2009; 4(7):1073–1082.

22. Choi Y, Sims GE, Murphy S, Miller JR, Chan AP. Predicting the functional effect of amino acid substitutions and indels. Plos One 2012; 7(10): e46688.

23. Adzhubei IA, Schmidt S, Peshkin L, Ramensky VE, Gerasimova A, Bork P, Kondrashov AS, Sunyaev SR. A method and server for predicting damaging missense mutations. Nat Methods 2010; 7(4):248–249.

24. Sherry ST, Ward MH, Kholodov M, Baker J, Phan L, Smigielski EM, et al. dbSNP: the NCBI database of genetic variation. Nucleic Acids Res 2001; 29(1):308–311.

25. New SNP Attributes 2018. Available at https://www.ncbi.nlm.nih.gov/projects/SNP/docs/rs_attributes.html.

26. Sim NL, Kumar P, Hu J, Henikoff S, Schneider G, Ng PC. SIFT web server: predicting effects of amino acid substitutions on proteins. Nucleic Acids Res 2012; 40: 542–547.

27. Ng PC, Henikoff S. Predicting deleterious amino acid substitutions. Genome Res 2001; 11(5):863–874.

28. Ng PC, Henikoff S. SIFT: Predicting amino acid changes that affect protein function. Nucleic Acids Res 2003; 31(13):3812–3814.

29. Yongwook C, Chan AP. PROVEAN web server: a tool to predict the functional effect of amino acid substitutions and indels. Bioinformatics 2015; 31(16):2745–2747.

30. Choi Y. 2012. A fast computation of pairwise sequence alignment scores between a protein and a set of single-locus variants of another protein. In: Proceedings of the ACM Conference on Bioinformatics, Computational Biology and Biomedicine; 2012 Oct 7; Orlando, Florida: ACM; p. 414–417.

31. Choi Y, Chan AP. PROVEAN web server: a tool to predict the functional effect of amino acid substitutions and indels. Bioinformatics Oxford England 2015; 31(16):2745–2747.

32. Sunyaev SR, Eisenhaber F, Rodchenkov IV, Eisenhaber B, Tumanyan VG, Kuznetsov EN. PSIC: profile extraction from sequence alignments with position-specific counts of independent observations. Protein Eng 1999; 12(5):387–394.

33. MacQueen J. Some methods for classification and analysis of multivariate observations. Proceedings of the Fifth Berkeley Symposium on Mathematical Statistics and Probability. 1967; 1(14):281–297.

34. Ester M, Kriegel HP, Sander J, Xu X. A density-based algorithm for discovering clusters in large spatial databases with noise. E Simoudis, J Han, MF Usama (Ed.). In: Proceedings of the Second International Conference on Knowledge Discovery and Data Mining (KDD-96) AAAI Press; 1996. p. 226–231.

35. Hochbaum DS, Shmoys DB. A best possible heuristic for the k-center problem. Math Oper Res 1985; 10(2):180–184.

36. Dasgupta S. Performance Guarantees for Hierarchical Clustering, In: 15th Annual Conference on Computational Learning Theory; 2002. p. 351–363.

37. Lakhani J, Khunteta A, Chowdhary A, Harwani D. Auto-Evolving Clusters based on Rejection and Migration. In: Proceedings of the International Conference on Advances in Information Communication Technology & Computing (AICTC 16). New York:Association for Computing Machinery; 2016. 98:1–6.

38. Alshatwi AA, Hasan TN, Syed NA, Shafi G, Grace BL. Identification of functional SNPs in BARD1 gene and in silico analysis of damaging snps: based on data procured from dbSNP database. Plos One 2012; 7(10): e43939.

39. Johnson N, Fletcher O, Palles C, Rudd M, Webb E, Sellick G,et al. Counting potentially functional variants in BRCA1, BRCA2 and ATM predicts breast cancer susceptibility. Hum Mol Genet 2007; 16(9):1051–1057.

40. Bullock AN, Henckel J, DeDecker BS, Johnson CM, Nikolova PV, Proctor MR, et al. Thermodynamic stability of wild-type and mutant p53 core domain. Proceedings of the National Academy of Sciences of United States of America 1997; 94(26):14338–14342.

41. Frazer KA, Murray SS, Schork NJ, Topol EJ. Human genetic variation and its contribution to complex traits. Nat Rev Genet 2009; 10(4): 241–251.

