## Supplementary Data Appendix A and B CA27.29 for "*In silico* analysis and cluster validation of potential breast cancer nsSNPs of serological tumor marker CA27.29"

### Appendix A Summary of deleterious nsSNPs predicted by SIFT tool

| SNP ID | Variation in Nucleotide | | | Variation in Amino Acid | | | Tolerance Value Cutoff - 0.05 |
|---|---|---|---|---|---|---|---|
| | Accession Number | Allele variation | Location of Allele Variation | Accession Number | Residue variation | Location of Residue Variation | |
| rs1611770 | NM_001018016.2 | G\|A | 448 | NP_001018016.1 | V\|M | 126 | 0.026 |
| | NM_001018017.2 | | 421 | NP_001018017.1 | | 117 | 0.031 |
| | NM_001204285.1 | | 1081 | NP_001191214.1 | | 337 | 0.013 |
| | NM_001204286.1 | | 1108 | NP_001191215.1 | | 346 | 0.012 |
| | NM_001204287.1 | | 502 | NP_001191216.1 | | 144 | 0.026 |
| | NM_001204288.1 | | 448 | NP_001191217.1 | | 126 | 0.021 |
| | NM_001204289.1 | | 370 | NP_001191218.1 | | 100 | 0.029 |
| | NM_001204290.1 | | 307 | NP_001191219.1 | | 79 | 0.015 |
| | NM_001204291.1 | | 397 | NP_001191220.1 | | 103 | 0.025 |
| | NM_001204292.1 | | 373 | NP_001191221.1 | | 101 | 0.026 |
| | NM_001204293.1 | | 475 | NP_001191222.1 | | 135 | 0.018 |
| | NM_001204294.1 | | 346 | NP_001191223.1 | | 92 | 0.032 |
| | NM-002456.5 | | 475 | NP_002449.4 | | 135 | 0.024 |
| | NM_001204285.1 | G\|T | 1081 | NP_001191214.1 | V\|L | 337 | 0.030 |
| | NM_001204286.1 | | 1108 | NP_001191215.1 | | 346 | 0.026 |
| | NM_001204287.1 | | 502 | NP_001191216.1 | | 144 | 0.046 |
| | NM_001204288.1 | | 448 | NP_001191217.1 | | 126 | 0.044 |
| | NM_001204293.1 | | 475 | NP_001191222.1 | | 135 | 0.044 |
| rs11465205 | NM_001204285.1 | G\|T | 658 | NP_001191214.1 | G\|C | 196 | 0.006 |
| | NM_001204286.1 | | 685 | NP_001191215.1 | | 205 | 0.005 |
| rs11465207 | NM_001037856.1 | G\|A | 241 | NP_001037856.1 | S\|N | 57 | 0.019 |
| | NM_001204285.1 | | 901 | NP_001191214.1 | | 277 | **0.000** |
| | NM_001204286.1 | | 928 | NP_001191215.1 | | 286 | 0.002 |
| | NM_001204287.1 | | 322 | NP_001191216.1 | | 84 | **0.000** |
| | NM_001204288.1 | | 268 | NP_001191217.1 | | 66 | 0.002 |
| | NM_001204293.1 | | 295 | NP_001191222.1 | | 75 | **0.000** |
| | NM_002456.5 | | 295 | NP_002447.4 | | 75 | **0.000** |
| rs35819649 | NM_001044391.2 | T\|A | 241 | NP_001037856.1 | S\|T | 57 | 0.019 |
| | NM_001204285.1 | | 901 | NP_001191214.1 | | 277 | **0.000** |
| | NM_001204286.1 | | 928 | NP_001191215.1 | | 286 | 0.002 |
| | NM_001204287.1 | | 322 | NP_001191216.1 | | 84 | **0.000** |
| | NM_001204288.1 | | 268 | NP_001191217.1 | | 66 | 0.002 |
| | NM_001204293.1 | | 295 | NP_001191222.1 | | 75 | **0.000** |
| | NM_002456.5 | | 295 | NP_002447.4 | | 75 | **0.000** |
| rs139437006 | NM_001018016.2 | C\|A | 134 | NP_001018016.1 | T\|K | 31 | **0.000** |
| | NM_001018017.2 | | 137 | NP_001018017.2 | | 22 | 0.002 |
| | NM_001044390.2 | | 137 | NP_001037855.1 | | 22 | **0.000** |
| | NM_001044391.2 | | 137 | NP_001037856.1 | | 22 | **0.000** |
| | NM_001044392.2 | | 164 | NP_001037857.1 | | 31 | **0.000** |
| | NM_001044393.2 | | 137 | NP_001037858.1 | | 22 | **0.000** |
| | NM_001204285.1 | | 137 | NP_001191214.1 | | 22 | 0.009 |
| | NM_001204286.1 | | 164 | NP_001191215.1 | | 31 | 0.002 |
| | NM_001204287.1 | | 164 | NP_001191216.1 | | 31 | 0.001 |
| | NM_001204288.1 | | 164 | NP_001191217.1 | | 31 | **0.000** |
| | NM_001204289.1 | | 164 | NP_001191218.1 | | 31 | **0.000** |
| | NM_001204291.1 | | 164 | NP_001191220.1 | | 31 | **0.000** |
| | NM_001204292.1 | | 164 | NP_001191221.1 | | 31 | **0.000** |
| | NM_001204293.1 | | 164 | NP_001191222.1 | | 22 | 0.001 |
| | NM_001204295.1 | | 164 | NP_001191224.1 | | 31 | **0.000** |
| | NM_001204296.1 | | 164 | NP_001191225.1 | | 31 | **0.000** |
| | NM_001204294.1 | | 164 | NP_001191226.1 | | 31 | **0.000** |
| | NM_002456.5 | | 137 | NP_002447.4 | | 22 | 0.001 |
| | NM_001018016.2 | C\|T | 134 | NP_001018016.1 | T\|M | 31 | **0.000** |

| | | | | | | | |
|---|---|---|---|---|---|---|---|
| | NM_001018017.2 | | 137 | NP_001018017.2 | | 22 | 0.002 |
| | NM_001044390.2 | | 137 | NP_001037855.1 | | 22 | 0.001 |
| | NM_001044391.2 | | 137 | NP_001037856.1 | | 22 | **0.000** |
| | NM_001044392.2 | | 164 | NP_001037857.1 | | 31 | **0.000** |
| | NM_001044393.2 | | 137 | NP_001037858.1 | | 22 | **0.000** |
| | NM_001204285.1 | | 137 | NP_001191214.1 | | 22 | 0.049 |
| | NM_001204286.1 | | 164 | NP_001191215.1 | | 31 | 0.001 |
| | NM_001204287.1 | | 164 | NP_001191216.1 | | 31 | **0.000** |
| | NM_001204288.1 | | 164 | NP_001191217.1 | | 31 | **0.000** |
| | NM_001204289.1 | | 164 | NP_001191218.1 | | 31 | **0.000** |
| | NM_001204291.1 | | 164 | NP_001191220.1 | | 31 | **0.000** |
| | NM_001204292.1 | | 164 | NP_001191221.1 | | 31 | **0.000** |
| | NM_001204293.1 | | 164 | NP_001191222.1 | | 22 | 0.001 |
| | NM_001204295.1 | | 164 | NP_001191224.1 | | 31 | **0.000** |
| | NM_001204296.1 | | 164 | NP_001191225.1 | | 31 | **0.000** |
| | NM_001204294.1 | | 164 | NP_001191226.1 | | 31 | **0.000** |
| | NM_002456.5 | | 137 | NP_002447.4 | | 22 | 0.001 |
| rs140871200 | NM_001044393.2 | G\|A | 305 | NP_001037858.1 | G\|G | 78 | **0.000** |
| rs141460657 | NM_001044393.2 | G\|T | 403 | NP_001037858.1 | G\|G | 111 | 0.003 |
| rs144273480 | NM_001018016.1 | G\|T | 445 | NP_001018016.1 | D\|Y | 125 | 0.009 |
| | NM_001018017.2 | | 418 | NP_001018017.1 | | 116 | 0.012 |
| | NM_001204285.1 | | 1078 | NP_001191214.1 | | 336 | 0.010 |
| | NM_001204286.1 | | 1105 | NP_001191215.1 | | 345 | 0.012 |
| | NM_001204287.1 | | 499 | NP_001191216.1 | | 143 | 0.018 |
| | NM_001204288.1 | | 445 | NP_001191217.1 | | 125 | 0.004 |
| | NM_001204289.1 | | 367 | NP_001191218.1 | | 99 | 0.028 |
| | NM_001204290.1 | | 304 | NP_001191219.1 | | 78 | 0.041 |
| | NM_001204291.1 | | 376 | NP_001191220.1 | | 102 | 0.007 |
| | NM_001204292.1 | | 370 | NP_001191221.1 | | 100 | 0.022 |
| | NM_001204293.1 | | 472 | NP_001191222.1 | | 134 | 0.006 |
| | NM_001204294.1 | | 343 | NP_001191223.1 | | 91 | 0.027 |
| | NM_002456.5 | | 472 | NP_002447.4 | | 134 | 0.011 |
| rs145108039 | NM_001018016.2 | G\|T | 182 | NP_001018016.1 | S\|I | 37 | 0.001 |
| | NM_001018017.2 | | 155 | NP_001018017.1 | | 28 | 0.001 |
| | NM_001044390.2 | | 155 | NP_001037855.1 | | 28 | 0.001 |
| | NM_001044391.2 | | 155 | NP_001037856.1 | | 28 | 0.001 |
| | NM_001044392.1 | | 182 | NP_001037857.1 | | 37 | 0.001 |
| | NM_001044393.2 | | 155 | NP_001037858.1 | | 28 | 0.001 |
| | NM_001204285.1 | | 155 | NP_001191214.1 | | 28 | 0.003 |
| | NM_001204286.1 | | 182 | NP_001191215.1 | | 37 | 0.001 |
| | NM_001204287.1 | | 182 | NP_001191216.1 | | 37 | 0.001 |
| | NM_001204288.1 | | 182 | NP_001191217.1 | | 37 | 0.001 |
| | NM_001204289.1 | | 182 | NP_001191218.1 | | 37 | 0.001 |
| | NM_001204291.1 | | 182 | NP_001191220.1 | | 37 | **0.000** |
| | NM_001204292.1 | | 182 | NP_001191221.1 | | 37 | **0.000** |
| | NM_001204293.1 | | 155 | NP_001191222.1 | | 28 | 0.001 |
| | NM_001204294.1 | | 155 | NP_001191223.1 | | 28 | 0.001 |
| | NM_001204295.1 | | 110 | NP_001191224.1 | | 28 | 0.001 |
| rs145224844 | NM_001018016.2 | G\|T | 389 | NP_001018016.1 | G\|V | 106 | 0.001 |
| | NM_001018017.2 | | 362 | NP_001018017.1 | | 97 | 0.001 |
| | NM_001204285.1 | | 1022 | NP_001191214.1 | | 317 | 0.001 |
| | NM_001204286.1 | | 1049 | NP_001191215.1 | | 326 | 0.001 |
| | NM_001204287.1 | | 443 | NP_001191216.1 | | 124 | 0.001 |
| | NM_001204288.1 | | 389 | NP_001191217.1 | | 106 | 0.001 |
| | NM_001204289.1 | | 311 | NP_001191218.1 | | 80 | 0.001 |
| | NM_001204290.1 | | 248 | NP_001191219.1 | | 59 | 0.001 |
| | NM_001204291.1 | | 320 | NP_001191220.1 | | 83 | 0.001 |
| | NM_001204292.1 | | 314 | NP_001191221.1 | | 81 | 0.001 |
| | NM_001204293.1 | | 416 | NP_001191222.1 | | 115 | 0.001 |
| | NM_001204294.1 | | 287 | NP_001191223.1 | | 72 | 0.001 |

| | | | | | | | |
|---|---|---|---|---|---|---|---|
| | NM_002456.5 | | 416 | NP_002447.4 | | 115 | 0.001 |
| rs145667707 | NM_001018016.2 | C\|T | 185 | NP_001018016.1 | S\|F | 38 | 0.001 |
| | NM_001018017.2 | | 158 | NP_001018017.1 | | 29 | 0.001 |
| | NM_001044390.2 | | 158 | NP_001037855.1 | | 29 | 0.001 |
| | NM_001044391.2 | | 158 | NP_001037856.1 | | 29 | 0.001 |
| | NM_001044392.2 | | 185 | NP_001037857.1 | | 38 | 0.001 |
| | NM_001044393.2 | | 158 | NP_001037858.1 | | 29 | 0.001 |
| | NM_001204285.1 | | 158 | NP_001191214.1 | | 29 | 0.001 |
| | NM_001204286.1 | | 185 | NP_001191215.1 | | 38 | **0.000** |
| | NM_001204287.1 | | 185 | NP_001191216.1 | | 38 | **0.000** |
| | NM_001204288.1 | | 185 | NP_001191217.1 | | 38 | 0.001 |
| | NM_001204289.1 | | 185 | NP_001191218.1 | | 38 | 0.001 |
| | NM_001204291.1 | | 185 | NP_001191220.1 | | 38 | **0.000** |
| | NM_001204292.1 | | 185 | NP_001191221.1 | | 38 | **0.000** |
| | NM_001204293.1 | | 158 | NP_001191222.1 | | 29 | 0.001 |
| | NM_001204294.1 | | 158 | NP_001191223.1 | | 29 | **0.000** |
| | NM_001204295.1 | | 185 | NP_001191224.1 | | 38 | 0.001 |
| | NM_001204296.1 | | 185 | NP_001191225.1 | | 38 | **0.000** |
| | NM_001204297.1 | | 185 | NP_001191226.1 | | 38 | 0.001 |
| | NM_002456.5 | | 158 | NP_002447.4 | | 29 | 0.003 |
| rs145691584 | NM_001018016.2 | C\|A | 240 | NP_001018016.1 | S\|R | 56 | 0.001 |
| | NM_001018017.2 | | 213 | NP_001018017.1 | | 47 | 0.004 |
| | NM_001044390.2 | | 213 | NP_001037855.1 | | 47 | 0.001 |
| | NM_001044391.2 | | 213 | NP_001037856.1 | | 47 | 0.001 |
| | NM_001044392.2 | | 240 | NP_001037857.1 | | 56 | 0.001 |
| | NM_001044393.2 | | 213 | NP_001037858.1 | | 47 | 0.001 |
| | NM_001204286.1 | | 240 | NP_001191215.1 | | 56 | 0.004 |
| | NM_001204287.1 | | 240 | NP_001191216.1 | | 56 | 0.010 |
| | NM_001204288.1 | | 240 | NP_001191217.1 | | 56 | 0.001 |
| | NM_001204289.1 | | 240 | NP_001191218.1 | | 56 | 0.003 |
| | NM_001204290.1 | | 177 | NP_001191219.1 | | 35 | 0.001 |
| | NM_001204291.1 | | 240 | NP_001191220.1 | | 56 | 0.002 |
| | NM_001204292.1 | | 240 | NP_001191221.1 | | 56 | 0.002 |
| | NM_001204293.1 | | 213 | NP_001191223.1 | | 47 | 0.003 |
| | NM_001204294.1 | | 213 | NP_001191223.1 | | 47 | 0.003 |
| | NM_001204295.1 | | 240 | NP_001191224.1 | | 56 | 0.002 |
| | NM_001204296.1 | | 240 | NP_001191225.1 | | 56 | 0.001 |
| | NM_001204297.1 | | 240 | NP_001191226.1 | | 56 | 0.004 |
| | NM_002456.5 | | 213 | NP_002447.4 | | 47 | 0.015 |
| rs146141676 | NM_001018016.2 | C\|T | 503 | NP_001018016.1 | T\|M | 144 | 0.002 |
| | NM_001018017.2 | | 476 | NP_001018017.1 | | 135 | 0.002 |
| | NM_001204285.1 | | 1136 | NP_001191214.1 | | 355 | 0.002 |
| | NM_001204286.1 | | 1163 | NP_001191215.1 | | 364 | 0.002 |
| | NM_001204287.1 | | 557 | NP_001191216.1 | | 162 | 0.001 |
| | NM_001204288.1 | | 503 | NP_001191217.1 | | 144 | 0.001 |
| | NM_001204289.1 | | 425 | NP_001191218.1 | | 118 | 0.004 |
| | NM_001204290.1 | | 362 | NP_001191219.1 | | 97 | 0.006 |
| | NM_001204291.1 | | 434 | NP_001191220.1 | | 121 | 0.002 |
| | NM_001204292.1 | | 428 | NP_001191221.1 | | 119 | 0.003 |
| | NM_001204293.1 | | 530 | NP_001191222.1 | | 153 | 0.004 |
| | NM_001204294.1 | | 401 | NP_001191223.1 | | 110 | 0.003 |
| | NM_002456.5 | | 530 | NP_002447.4 | | 153 | 0.001 |
| rs146950322 | NM_001018016.2 | C\|T | 280 | NP_001018016.1 | P\|S | 70 | **0.000** |
| | NM_001018017.2 | | 253 | NP_001018017.1 | | 61 | **0.000** |
| | NM_001204285.1 | | 913 | NP_001191214.1 | | 281 | **0.000** |
| | NM_001204286.1 | | 940 | NP_001191215.1 | | 290 | **0.000** |
| | NM_001204287.1 | | 334 | NP_001191216.1 | | 88 | **0.000** |
| | NM_001204288.1 | | 280 | NP_001191217.1 | | 70 | **0.000** |
| | NM_001204293.1 | | 307 | NP_001191222.1 | | 79 | **0.000** |
| | NM_002456.5 | | 307 | NP_002447.4 | | 79 | **0.000** |

| | | | | | | | |
|---|---|---|---|---|---|---|---|
| rs148332231 | NM_001018016.2 | A\|G | 785 | NP_001018016.1 | Y\|C | 238 | **0.000** |
| | NM_001044390.2 | | 602 | NP_001037855.1 | | 177 | **0.000** |
| | NM_001044391.2 | | 443 | NP_001037856.1 | | 124 | **0.000** |
| | NM_001044392.2 | | 470 | NP_001037857.1 | | 133 | **0.000** |
| | NM_001204285.1 | | 1418 | NP_001191214.1 | | 449 | **0.000** |
| | NM_001204286.1 | | 458 | NP_001191215.1 | | 458 | **0.000** |
| | NM_001204287.1 | | 839 | NP_001191216.1 | | 256 | **0.000** |
| | NM_001204289.1 | | 707 | NP_001191218.1 | | 212 | **0.000** |
| | NM_001204290.1 | | 644 | NP_001191219.1 | | 191 | **0.000** |
| | NM_001204291.1 | | 716 | NP_001191220.1 | | 215 | **0.000** |
| | NM_001204292.1 | | 710 | NP_001191221.1 | | 213 | **0.000** |
| | NM_001204294.1 | | 683 | NP_001191223.1 | | 204 | **0.000** |
| | NM_001204295.1 | | 560 | NP_001191224.1 | | 163 | **0.000** |
| | NM_001204296.1 | | 629 | NP_001191225.1 | | 186 | **0.000** |
| | NM_001204297.1 | | 524 | NP_001191226.1 | | 151 | **0.000** |
| rs149173724 | NM_001204286.1 | G\|T | 298 | NP_001191215.1 | G\|C | 76 | 0.014 |
| rs183700327 | NM_001204286.1 | A\|G | 1289 | NP_001191215.1 | Y\|C | 406 | 0.046 |
| rs191544901 | NM_001018016.2 | G\|A | 776 | NP_001018016.1 | R\|H | 235 | 0.016 |
| | NM_001018017.2 | | 749 | NP_001018017.1 | | 226 | 0.017 |
| | NM_001044390.2 | | 593 | NP_001031855.1 | | 174 | 0.014 |
| | NM_001044391.2 | | 434 | NP_001037856.1 | | 121 | 0.013 |
| | NM_001044392.2 | | 461 | NP_001037857.1 | | 130 | 0.017 |
| | NM_001204285.1 | | 1049 | NP_001191214.1 | | 446 | 0.029 |
| | NM_001204286.1 | | 1436 | NP_001191215.1 | | 455 | 0.018 |
| | NM_001204287.1 | | 830 | NP_001191216.1 | | 253 | 0.008 |
| | NM_001204289.1 | | 698 | NP_001191218.1 | | 209 | 0.008 |
| | NM_001204290.1 | | 635 | NP_001191219.1 | | 188 | 0.009 |
| | NM_001204291.1 | | 707 | NP_001191220.1 | | 212 | 0.011 |
| | NM_001204292.1 | | 701 | NP_001191221.1 | | 701 | 0.010 |
| | NM_001204294.1 | | 674 | NP_001191223.1 | | 201 | 0.015 |
| | NM_001204295.1 | | 551 | NP_001191224.1 | | 160 | 0.012 |
| | NM_001204296.1 | | 620 | NP_001191225.1 | | 183 | 0.010 |
| | NM_001204297.1 | | 515 | NP_001191226.1 | | 148 | 0.013 |
| | NM_002456.5 | | 803 | NP_002447.4 | | 244 | 0.022 |
| | NM_001018016.2 | G\|T | 776 | NP_001018016.1 | R\|L | 235 | 0.001 |
| | NM_001018017.2 | | 749 | NP_001018017.1 | | 226 | 0.001 |
| | NM_001044390.2 | | 593 | NP_001031855.1 | | 174 | 0.001 |
| | NM_001044391.2 | | 434 | NP_001037856.1 | | 121 | 0.001 |
| | NM_001044392.2 | | 461 | NP_001037857.1 | | 130 | 0.001 |
| | NM_001204285.1 | | 1049 | NP_001191214.1 | | 446 | 0.001 |
| | NM_001204286.1 | | 1436 | NP_001191215.1 | | 455 | 0.001 |
| | NM_001204287.1 | | 830 | NP_001191216.1 | | 253 | 0.001 |
| | NM_001204289.1 | | 698 | NP_001191218.1 | | 209 | 0.001 |
| | NM_001204290.1 | | 635 | NP_001191219.1 | | 188 | 0.001 |
| | NM_001204291.1 | | 707 | NP_001191220.1 | | 212 | 0.001 |
| | NM_001204292.1 | | 701 | NP_001191221.1 | | 701 | 0.001 |
| | NM_001204294.1 | | 674 | NP_001191223.1 | | 201 | 0.001 |
| | NM_001204295.1 | | 551 | NP_001191224.1 | | 160 | 0.001 |
| | NM_001204296.1 | | 620 | NP_001191225.1 | | 183 | 0.001 |
| | NM_001204297.1 | | 515 | NP_001191226.1 | | 148 | 0.001 |
| | NM_002456.5 | | 803 | NP_002447.4 | | 244 | 0.001 |

**Appendix B Summary of deleterious nsSNPs predicted by PROVEAN tool**

| SNP ID | Variation in Nucleotide | | | Variation in Amino Acid | | | Tolerance Value Cutoff |
|---|---|---|---|---|---|---|---|
| | Accession Number | Allele variation | Location of Allele Variation | Accession Number | Residue variation | Location of Residue Variation | |
| rs139437006 | NM_001018016.2 | C\|A | 134 | NP_001018016.1 | T\|K | 31 | **-2.92** |
| | NM_001018017.2 | | 137 | NP_001018017.2 | | 22 | -2.78 |
| | NM_001044391.2 | | 137 | NP_001037856.1 | | 22 | **-2.61** |
| | NM_001044392.2 | | 164 | NP_001037857.1 | | 31 | **-3.15** |
| | NM_001044393.2 | | 137 | NP_001037858.1 | | 22 | **-5.39** |
| | NM_001204287.1 | | 164 | NP_001191216.1 | | 31 | -2.93 |
| | NM_001204288.1 | | 164 | NP_001191217.1 | | 31 | **-3.55** |
| | NM_001204289.1 | | 164 | NP_001191218.1 | | 31 | **-2.88** |
| | NM_001204291.1 | | 164 | NP_001191220.1 | | 31 | **-3.02** |
| | NM_001204292.1 | | 164 | NP_001191221.1 | | 31 | **-3.29** |
| | NM_001204294.1 | | 137 | NP_001191223.1 | | 31 | -2.92 |
| | NM_001204296.1 | | 164 | NP_001191225.1 | | 31 | **-3.38** |
| | NM_001204294.1 | | 164 | NP_001191226.1 | | 31 | **-2.86** |
| | NM_001018016.2 | C\|T | 134 | NP_001018016.1 | T\|M | 31 | **-2.99** |
| | NM_001018017.2 | | 137 | NP_001018017.2 | | 22 | **-2.97** |
| | NM_001044391.2 | | 137 | NP_001037856.1 | | 22 | **-2.72** |
| | NM_001044392.2 | | 164 | NP_001037857.1 | | 31 | **-3.22** |
| | NM_001044393.2 | | 137 | NP_001037858.1 | | 22 | **-5.39** |
| | NM_001204286.1 | | 164 | NP_001191215.1 | | 31 | -2.66 |
| | NM_001204287.1 | | 164 | NP_001191216.1 | | 31 | **-3.04** |
| | NM_001204288.1 | | 164 | NP_001191217.1 | | 31 | **-3.57** |
| | NM_001204289.1 | | 164 | NP_001191218.1 | | 31 | **-3.01** |
| | NM_001204291.1 | | 164 | NP_001191220.1 | | 31 | **-3.02** |
| | NM_001204292.1 | | 164 | NP_001191221.1 | | 31 | **-3.30** |
| | NM_001204294.1 | | 137 | NP_001191223.1 | | 31 | -2.91 |
| | NM_001204296.1 | | 164 | NP_001191225.1 | | 31 | **-3.48** |
| | NM_001204294.1 | | 164 | NP_001191226.1 | | 31 | **-3.04** |
| rs144273480 | NM_001018016.1 | G\|T | 445 | NP_001018016.1 | D\|Y | 125 | -2.54 |
| | NM_001018017.2 | | 418 | NP_001018017.1 | | 116 | -2.54 |
| | NM_001204285.1 | | 1078 | NP_001191214.1 | | 336 | -2.93 |
| | NM_001204286.1 | | 1105 | NP_001191215.1 | | 345 | -2.94 |
| | NM_001204287.1 | | 499 | NP_001191216.1 | | 143 | -2.67 |
| | NM_001204288.1 | | 445 | NP_001191217.1 | | 125 | -2.62 |
| | NM_001204289.1 | | 367 | NP_001191218.1 | | 99 | -2.70 |
| | NM_001204290.1 | | 304 | NP_001191219.1 | | 78 | -2.57 |
| | NM_001204291.1 | | 376 | NP_001191220.1 | | 102 | -2.66 |
| | NM_001204292.1 | | 370 | NP_001191221.1 | | 100 | -2.76 |
| | NM_001204293.1 | | 472 | NP_001191222.1 | | 134 | -2.54 |
| | NM_001204294.1 | | 343 | NP_001191223.1 | | 91 | -2.86 |
| | NM_002456.5 | | 472 | NP_002447.4 | | 134 | -2.67 |
| rs145108039 | NM_001018016.2 | G\|T | 182 | NP_001018016.1 | S\|I | 37 | -3.07 |
| | NM_001018017.2 | | 155 | NP_001018017.1 | | 28 | -2.79 |
| | NM_001044391.2 | | 155 | NP_001037856.1 | | 28 | -2.56 |
| | NM_001044392.1 | | 182 | NP_001037857.1 | | 37 | -3.14 |
| | NM_001044393.2 | | 155 | NP_001037858.1 | | 28 | -5.56 |
| | NM_001204286.1 | | 182 | NP_001191215.1 | | 37 | -2.70 |
| | NM_001204287.1 | | 182 | NP_001191216.1 | | 37 | -3.17 |
| | NM_001204288.1 | | 182 | NP_001191217.1 | | 37 | -3.93 |
| | NM_001204289.1 | | 182 | NP_001191218.1 | | 37 | -2.85 |
| | NM_001204291.1 | | 182 | NP_001191220.1 | | 37 | **-2.79** |
| | NM_001204292.1 | | 182 | NP_001191221.1 | | 37 | **-2.99** |
| | NM_001204293.1 | | 155 | NP_001191222.1 | | 28 | -2.98 |
| | NM_001204294.1 | | 155 | NP_001191223.1 | | 28 | -2.98 |

| | | | | | | | |
|---|---|---|---|---|---|---|---|
| rs145224844 | NM_001018016.2 | G\|T | 389 | NP_001018016.1 | G\|V | 106 | -5.94 |
| | NM_001018017.2 | | 362 | NP_001018017.1 | | 97 | -5.94 |
| | NM_001204285.1 | | 1022 | NP_001191214.1 | | 317 | -6.31 |
| | NM_001204286.1 | | 1049 | NP_001191215.1 | | 326 | -6.32 |
| | NM_001204287.1 | | 443 | NP_001191216.1 | | 124 | -5.95 |
| | NM_001204288.1 | | 389 | NP_001191217.1 | | 106 | -5.41 |
| | NM_001204289.1 | | 311 | NP_001191218.1 | | 80 | -5.90 |
| | NM_001204290.1 | | 248 | NP_001191219.1 | | 59 | -5.89 |
| | NM_001204291.1 | | 320 | NP_001191220.1 | | 83 | -5.67 |
| | NM_001204292.1 | | 314 | NP_001191221.1 | | 81 | -5.81 |
| | NM_001204293.1 | | 416 | NP_001191222.1 | | 115 | -5.47 |
| | NM_001204294.1 | | 287 | NP_001191223.1 | | 72 | -5.89 |
| | NM_002456.5 | | 416 | NP_002447.4 | | 115 | -5.95 |
| rs145667707 | NM_001018016.2 | C\|T | 185 | NP_001018016.1 | S\|F | 38 | -3.67 |
| | NM_001018017.2 | | 158 | NP_001018017.1 | | 29 | -3.68 |
| | NM_001044391.2 | | 158 | NP_001037856.1 | | 29 | -3.55 |
| | NM_001044392.2 | | 185 | NP_001037857.1 | | 38 | -3.67 |
| | NM_001044393.2 | | 158 | NP_001037858.1 | | 29 | -5.64 |
| | NM_001204285.1 | | 158 | NP_001191214.1 | | 29 | -2.95 |
| | NM_001204286.1 | | 185 | NP_001191215.1 | | 38 | **-3.21** |
| | NM_001204287.1 | | 185 | NP_001191216.1 | | 38 | **-3.08** |
| | NM_001204288.1 | | 185 | NP_001191217.1 | | 38 | -4.23 |
| | NM_001204289.1 | | 185 | NP_001191218.1 | | 38 | -2.67 |
| | NM_001204291.1 | | 185 | NP_001191220.1 | | 38 | **-2.73** |
| | NM_001204292.1 | | 185 | NP_001191221.1 | | 38 | **-3.14** |
| | NM_001204293.1 | | 158 | NP_001191222.1 | | 29 | -3.40 |
| | NM_001204294.1 | | 158 | NP_001191223.1 | | 29 | **-3.00** |
| | NM_001204295.1 | | 185 | NP_001191224.1 | | 38 | -2.84 |
| | NM_001204297.1 | | 185 | NP_001191226.1 | | 38 | -3.47 |
| | NM_002456.5 | | 158 | NP_002447.4 | | 29 | -3.04 |
| rs145691584 | NM_001044393.2 | C\|A | 213 | NP_001037858.1 | S\|R | 47 | -4.80 |
| | NM_001204288.1 | | 240 | NP_001191217.1 | | 56 | -2.72 |
| | NM_001204297.1 | | 240 | NP_001191226.1 | | 56 | -2.70 |
| rs146950322 | NM_001018016.2 | C\|T | 280 | NP_001018016.1 | P\|S | 70 | **-6.09** |
| | NM_001018017.2 | | 253 | NP_001018017.1 | | 61 | **-6.26** |
| | NM_001044391.2 | | 253 | NP_001037856.1 | | 61 | **-3.54** |
| | NM_001044392.2 | | 280 | NP_001037857.1 | | 70 | **-4.62** |
| | NM_001204285.1 | | 913 | NP_001191214.1 | | 281 | **-6.02** |
| | NM_001204286.1 | | 940 | NP_001191215.1 | | 290 | **-6.08** |
| | NM_001204287.1 | | 334 | NP_001191216.1 | | 88 | **-6.47** |
| | NM_001204288.1 | | 280 | NP_001191217.1 | | 70 | **-6.22** |
| | NM_001204293.1 | | 307 | NP_001191222.1 | | 79 | **-6.19** |
| | NM_001204295.1 | | 280 | NP_001191224.1 | | 70 | **-4.76** |
| | NM_002456.5 | | 307 | NP_002447.4 | | 79 | **-6.50** |
| rs148332231 | NM_001018016.2 | A\|G | 785 | NP_001018016.1 | Y\|C | 238 | **-8.01** |
| | NM_001044390.2 | | 602 | NP_001037855.1 | | 177 | **-7.89** |
| | NM_001044391.2 | | 443 | NP_001037856.1 | | 124 | **-7.73** |
| | NM_001044392.2 | | 470 | NP_001037857.1 | | 133 | **-7.43** |
| | NM_001204285.1 | | 1418 | NP_001191214.1 | | 449 | **-7.94** |
| | NM_001204286.1 | | 458 | NP_001191215.1 | | 458 | **-7.94** |
| | NM_001204287.1 | | 839 | NP_001191216.1 | | 256 | **-7.96** |
| | NM_001204289.1 | | 707 | NP_001191218.1 | | 212 | **-8.01** |
| | NM_001204290.1 | | 644 | NP_001191219.1 | | 191 | **-7.99** |
| | NM_001204291.1 | | 716 | NP_001191220.1 | | 215 | **-8.01** |
| | NM_001204292.1 | | 710 | NP_001191221.1 | | 213 | **-8.01** |
| | NM_001204294.1 | | 683 | NP_001191223.1 | | 204 | **-7.85** |
| | NM_001204295.1 | | 560 | NP_001191224.1 | | 163 | **-7.92** |
| | NM_001204296.1 | | 629 | NP_001191225.1 | | 186 | **-7.37** |
| | NM_001204297.1 | | 524 | NP_001191226.1 | | 151 | **-7.96** |

| | | | | | | | |
|---|---|---|---|---|---|---|---|
| rs183700327 | NM_001018016.2 | A\|G | 629 | NP_001018016.1 | Y\|C | 186 | -6.65 |
| | NM_001018017.2 | | 602 | NP_001018017.1 | | 177 | -6.65 |
| | NM_001044390.2 | | 446 | NP_001037855.1 | | 125 | -6.46 |
| | NM_001204285.1 | | 1262 | NP_001191214.1 | | 397 | -6.48 |
| | NM_001204286.1 | | 1289 | NP_001191215.1 | | 406 | -6.49 |
| | NM_001204287.1 | | 683 | NP_001191216.1 | | 204 | -6.59 |
| | NM_001204289.1 | | 551 | NP_001191218.1 | | 551 | -6.62 |
| | NM_001204290.1 | | 488 | NP_001191219.1 | | 139 | -6.61 |
| | NM_001204291.1 | | 560 | NP_001191220.1 | | 163 | -6.63 |
| | NM_001204292.1 | | 554 | NP_001191221.1 | | 161 | -6.63 |
| | NM_001204294.1 | | 527 | NP_001191223.1 | | 152 | -6.57 |
| | NM_001204295.1 | | 404 | NP_001191224.1 | | 111 | -6.69 |
| | NM_001204296.1 | | 473 | NP_001191225.1 | | 134 | -6.48 |
| | NM_002456.5 | | 656 | NP_002447.4 | | 195 | -6.59 |
| rs191544901 | NM_001018016.2 | G\|A | 776 | NP_001018016.1 | R\|H | 235 | -4.45 |
| | NM_001018017.2 | | 749 | NP_001018017.1 | | 226 | -4.45 |
| | NM_001044390.2 | | 593 | NP_001031855.1 | | 174 | -4.40 |
| | NM_001044391.2 | | 434 | NP_001037856.1 | | 121 | -4.19 |
| | NM_001044392.2 | | 461 | NP_001037857.1 | | 130 | -3.90 |
| | NM_001204285.1 | | 1049 | NP_001191214.1 | | 446 | -4.41 |
| | NM_001204286.1 | | 1436 | NP_001191215.1 | | 455 | -4.41 |
| | NM_001204287.1 | | 830 | NP_001191216.1 | | 253 | -4.42 |
| | NM_001204289.1 | | 698 | NP_001191218.1 | | 209 | -4.45 |
| | NM_001204290.1 | | 635 | NP_001191219.1 | | 188 | -4.45 |
| | NM_001204291.1 | | 707 | NP_001191220.1 | | 212 | -4.45 |
| | NM_001204292.1 | | 701 | NP_001191221.1 | | 701 | -4.45 |
| | NM_001204294.1 | | 674 | NP_001191223.1 | | 201 | -4.45 |
| | NM_001204295.1 | | 551 | NP_001191224.1 | | 160 | -4.38 |
| | NM_001204296.1 | | 620 | NP_001191225.1 | | 183 | -4.42 |
| | NM_001204297.1 | | 515 | NP_001191226.1 | | 148 | -3.89 |
| | NM_002456.5 | | 803 | NP_002447.4 | | 244 | -4.42 |
| | NM_001018016.2 | G\|T | 776 | NP_001018016.1 | R\|L | 235 | -6.27 |
| | NM_001018017.2 | | 749 | NP_001018017.1 | | 226 | -6.27 |
| | NM_001044390.2 | | 593 | NP_001031855.1 | | 174 | -6.18 |
| | NM_001044391.2 | | 434 | NP_001037856.1 | | 121 | -5.97 |
| | NM_001044392.2 | | 461 | NP_001037857.1 | | 130 | -5.85 |
| | NM_001204285.1 | | 1049 | NP_001191214.1 | | 446 | -6.20 |
| | NM_001204286.1 | | 1436 | NP_001191215.1 | | 455 | -6.20 |
| | NM_001204287.1 | | 830 | NP_001191216.1 | | 253 | -6.22 |
| | NM_001204289.1 | | 698 | NP_001191218.1 | | 209 | -6.27 |
| | NM_001204290.1 | | 635 | NP_001191219.1 | | 188 | -6.26 |
| | NM_001204291.1 | | 707 | NP_001191220.1 | | 212 | -6.27 |
| | NM_001204292.1 | | 701 | NP_001191221.1 | | 701 | -6.27 |
| | NM_001204294.1 | | 674 | NP_001191223.1 | | 201 | -6.26 |
| | NM_001204295.1 | | 551 | NP_001191224.1 | | 160 | -6.22 |
| | NM_001204296.1 | | 620 | NP_001191225.1 | | 183 | -6.20 |
| | NM_001204297.1 | | 515 | NP_001191226.1 | | 148 | -5.69 |
| | NM_002456.5 | | 803 | NP_002447.4 | | 244 | -6.22 |
